# Speciation genes are more likely to have discordant gene trees

**DOI:** 10.1101/244822

**Authors:** Richard J. Wang, Matthew W. Hahn

## Abstract

Speciation genes are responsible for reproductive isolation between species. By directly participating in the process of speciation, the genealogies of isolating loci have been thought to more faithfully represent species trees. The unique properties of speciation genes may provide valuable evolutionary insights and help determine the true history of species divergence. Here, we formally analyze whether genealogies from loci participating in Dobzhansky-Muller (DM) incompatibilities are more likely to be concordant with the species tree under incomplete lineage sorting (ILS). Individual loci differ stochastically from the true history of divergence with a predictable frequency due to ILS, and these expectations—combined with the DM model of intrinsic reproductive isolation from epistatic interactions—can be used to examine the probability of concordance at isolating loci. Contrary to existing verbal models, we find that reproductively isolating loci that follow the DM model are often more likely to have discordant gene trees. These results are dependent on the pattern of isolation observed between three species, the time between speciation events, and the time since the last speciation event. Results supporting a higher probability of discordance are found for both derived-derived and derived-ancestral DM pairs, and regardless of whether incompatibilities are allowed or prohibited from segregating in the same population. Our overall results suggest that DM loci are unlikely to be especially useful for reconstructing species relationships, even in the presence of gene flow between incipient species, and may in fact be positively misleading.

## Introduction

Speciation proceeds from the evolution of reproductive isolation between populations. The study of reproductive isolation has advanced our understanding of the genetic basis of speciation, for which a common evolutionary model has become established. The Dobzhansky-Muller (DM) model describes how hybrid incompatibilities can arise as the result of epistasis between two or more loci that have diverged between populations (Bateson 1909; Dobzhansky 1937; Muller 1940). By having incompatible alleles for these loci arise in separate populations, the DM model allows reproductive isolation to evolve between populations without the appearance of reproductive failure within populations. A growing number of so-called ‘speciation genes’ that isolate species in accordance with the DM model have emerged from the genetic analysis of reproductive isolation in hybrids (e.g. Ting et al. 1998; Barbash et al. 2003; Presgraves et al. 2003; Bomblies et al. 2007; Mihola et al. 2009; Phadnis and Orr 2009; Barr and Fishman 2010; Lienard et al. 2016). Combinations of alleles from different species at these genes cause hybrid infertility or inviability.

The identification of speciation genes in multiple model systems has led to a search for the genetic, molecular, and evolutionary commonalities among them (Orr et al. 2004; Wu and Ting 2004; Presgraves 2010; Rieseberg and Blackman 2010; Nosil and Schluter 2011; Castillo and Barbash 2017). A major question is whether the genes leading to reproductive isolation differ from other genes in the genome. Various hypotheses have suggested that speciation genes are more likely to be the targets of adaptive evolution (Coyne and Orr 2004), more prone to interact with other genes (Guerrero et al. 2017), or more likely to be involved in genetic conflict (Bomblies et al. 2007; Phadnis and Orr 2009; Agren 2013).

The unique role of speciation genes in establishing species boundaries has also led to arguments asserting that these genes should be especially informative about species relationships (Ting et al. 2000; Rosenberg 2003; Zachos 2009; Maroja et al. 2009; Nosil and Schluter 2011; Cutter 2013). Such a property becomes useful when multiple species are separated by very short times between successive speciation events. In these cases, individual gene trees may have different topologies from one another and from the species tree (Maddison 1997). This phenomenon is not due to low power or sampling error, but represents a real difference in the genealogical history between loci, due to incomplete lineage sorting (ILS) or gene flow. With the high degree of discordance seen in many systems (e.g. Pollard et al. 2006; White et al. 2009; Jarvis et al. 2014; Pease et al. 2016), concordance between the topology of speciation genes and species trees would provide uniquely powerful insight into evolutionary histories. Verbal models have created the impression that speciation loci are biased towards concordance (Ting et al. 2000; Nosil and Schluter 2011; Cutter 2013), but no formal analysis of this idea has been carried out.

Here, we compare the expected genealogical history of loci involved in DMIs to the expected history of loci uninvolved in incompatibilities from the same genomes. The appreciation of discordance among gene trees has become acute with whole-genome sequence data, leading to multiple methods that incorporate ILS in the inference of species trees (e.g. Liu et al. 2009; Heled and Drummond 2010; Larget et al. 2010; Mirarab and Warnow 2015). However, gene tree discordance has received limited consideration in the context of DMIs. Loci involved in DMIs present additional challenges because of the epistatic nature of incompatibilities, which means that participating loci must act together to produce the incompatible phenotype. In addition, because both alleles involved in an incompatibility cannot segregate in the same population without leading to lower fitness in some individuals, the order in which mutations arise at each locus in a DMI matters.

We find that under a neutral model with incomplete lineage sorting, the stochastic processes of mutation and coalescence typically lead to higher rates of species tree discordance at hybrid incompatibility loci. We arrive at this counterintuitive result by examining the probability of ILS at loci participating in a canonical two-locus DMI. Our analysis considers four potential types of gene trees at a hypothetical incompatibility locus (Fig. 1). A key initial insight is recognizing the possibility that incompatible alleles can arise on discordant gene trees and still lead to reproductive isolation between pairs of species. Figure 2 shows how a DMI can arise from two loci, both with discordant gene trees, and isolate one or more species pairs. Since the expected branch lengths for each type of gene tree differ, mutations giving rise to incompatible alleles are not equally likely among the types of gene trees. We consider each combination of topologies for a pair of loci and calculate the probability of a DMI from differences in expected branch lengths. We find that loci participating in DMIs are typically more likely to have discordant gene trees, and that some patterns of isolation between species are more likely when loci are discordant.

**Figure 1.**
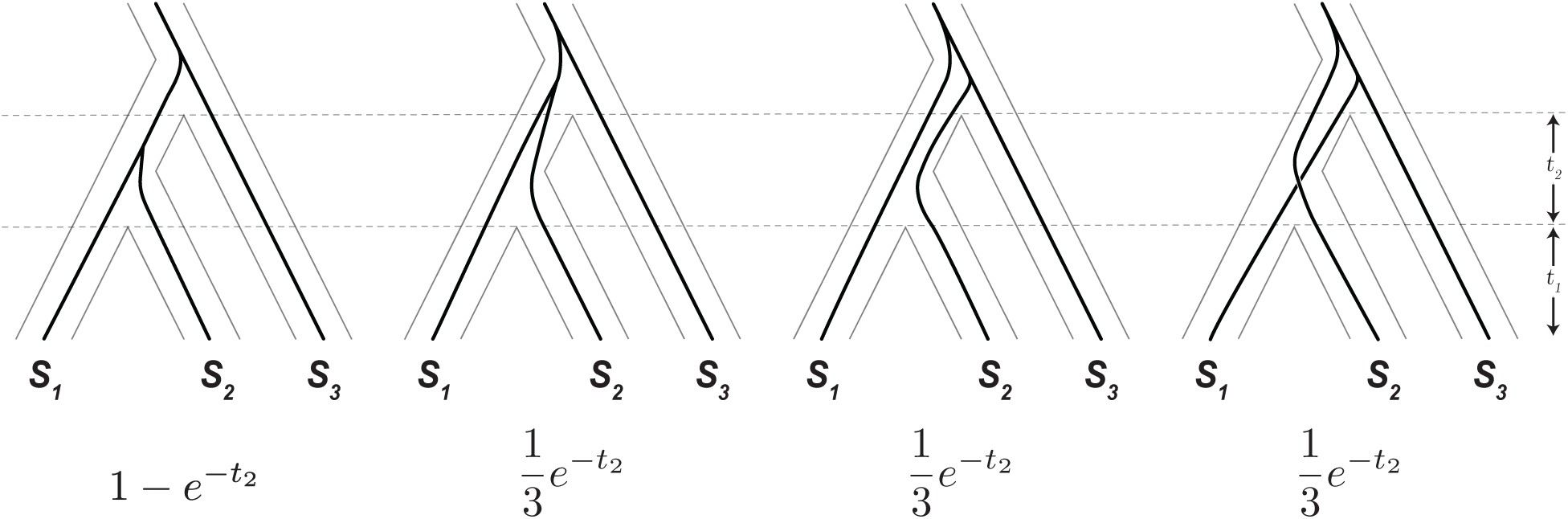
Four types of gene trees at a DMI locus and their expected frequencies. Two gene trees concordant with the species tree (left), and two that are discordant (right). Only the leftmost gene tree coalesces before the first speciation event. The labeled times, *t_1_* and *t_2_*, are the time from present to the first speciation event and the time between speciation events, respectively. Below each gene tree is the probability of its occurrence for a random locus in the genome under ILS alone.

**Figure 2.**
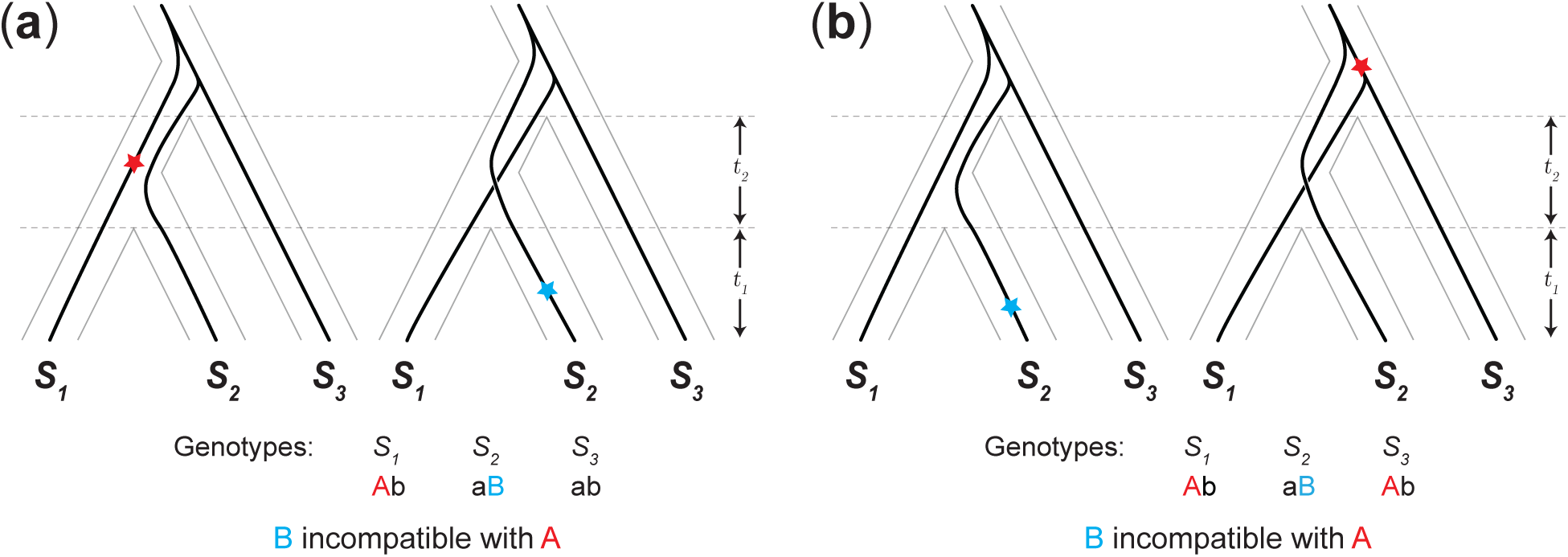
Incompatibility between one or more species pairs due to alleles from two loci that both have discordant gene trees, (*S_2_*, *S_3_*) *S_1_* and (*S_1_*, *S3*) *S_2_*. Red and blue stars mark the position of the first and second mutations, respectively, that produce incompatible alleles. The ancestral genotype for the two loci is denoted ‘ab’, with the mutations producing derived alleles ‘A’ and ‘B’. (a) Incompatibility between lineages *S_1_* and *S_2_*. (b) Incompatibility between lineages *S_1_* and *S_2_* as well as *S_3_* and *S_2_* from shared incompatible alleles.

## Results

### Preliminaries

Our genealogical model considers a single pairwise DMI in a three-species complex. DMIs are typically modeled as isolating two taxa, but depending on where an incompatible allele arises on a phylogenetic tree, a DMI can be shared among different species pairs (Moyle and Payseur 2009). The most straightforward way for this to occur is to have an interaction between two derived alleles (“derived-derived” incompatibilities; Orr 1995), where one of the derived alleles is shared between species, having arisen before their divergence (Fig. 2b). Two mutations inherited by the same lineage can also result in shared isolation, with the second derived allele being incompatible with the ancestral allele in other taxa (“derived-ancestral” incompatibilities; Orr 1995; Fig. S1). Generally, a DMI involving only two loci can produce six patterns of isolation among three taxa. The topology and branch lengths for the two most recently diverged taxa are interchangeable in many of the subsequent calculations, leaving four unique patterns of reproductive isolation (Figure 3). As we show below, these patterns of reproductive isolation are more often associated with particular types of gene trees.

**Figure 3.**
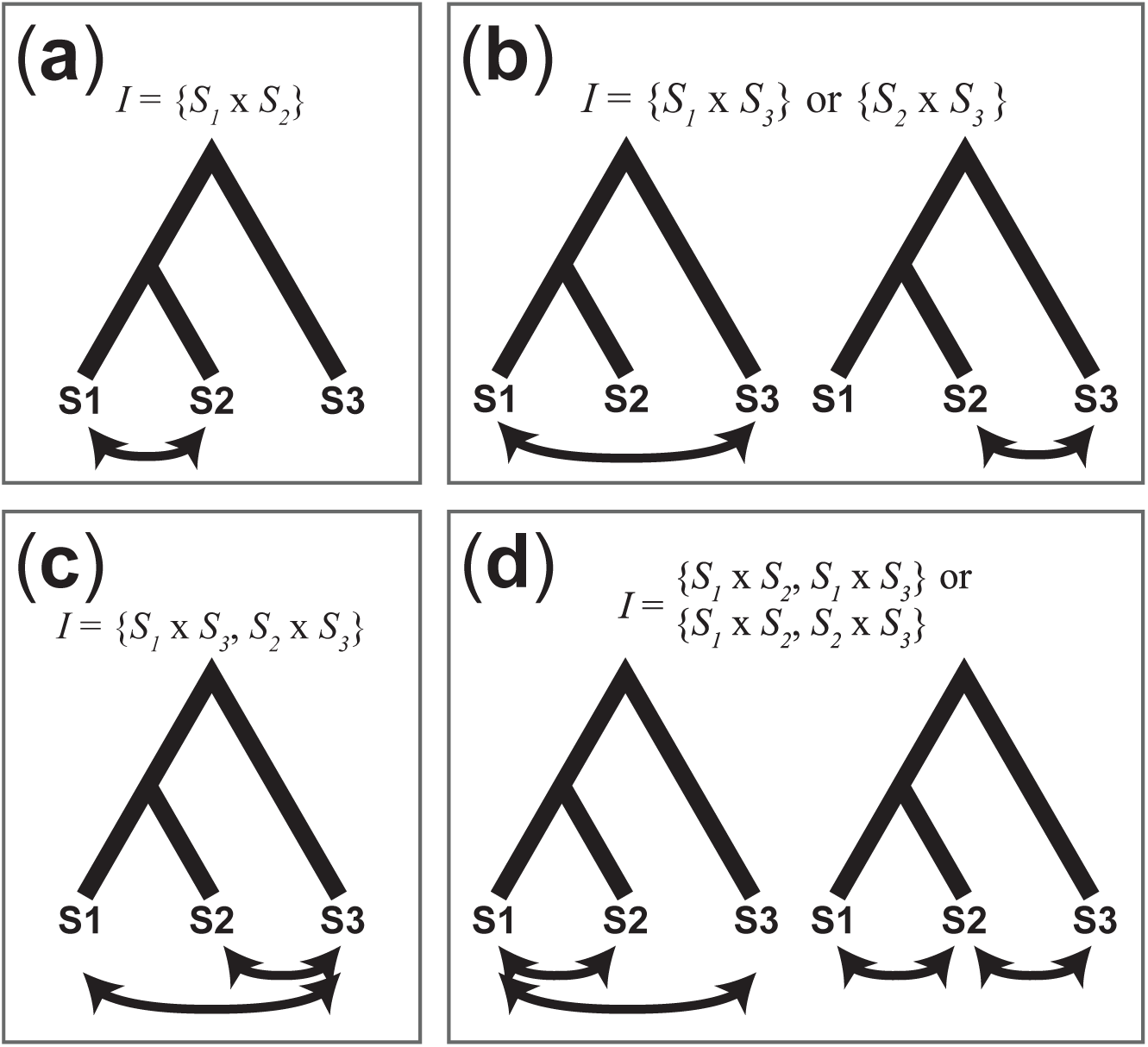
Four patterns of reproductive isolation. A single pairwise DMI can isolate a pair of species, as in panels (a) and (b), or two pairs of species, as in panels (c) and (d). For subsequent calculations, *S_1_* and *S_2_* are often interchangeable, leading us to group the two sets of relationships in (b) and (d).

We allow loci participating in an incompatibility to be discordant with the species tree only through incomplete lineage sorting. Specifically, a DMI locus can have one of four potential types of gene trees (Fig. 1). While there are only three potential topologies for three species, we divide concordant gene trees into those that coalesce in the ancestral population and those that coalesce between the speciation events (i.e. are lineage-sorted). Discordant gene trees must coalesce in the ancestral population of all three species. For a single locus, each of the three ancestrally coalescing gene trees are equally likely, at 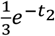, where *t_2_* is the inter-speciation time (Hudson 1983; Nei 1986). This leaves the probability of a concordant, lineage-sorted gene tree at 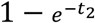. For two independent loci, the joint probability of any particular pair of genealogies is a product of the probabilities above. However, this is not the case for two loci participating in a DMI.

#### Calculating concordance at a DMI locus

For a given pattern of reproductive isolation, certain pairs of gene trees are more likely to give rise to incompatible alleles. Loci participating in a DMI must have experienced mutations on the appropriate branches to form the corresponding pattern of isolation. As an obvious example, a mutation specific to the *S_3_* lineage, on any of the gene trees shown in Figure 1, cannot create an incompatibility that isolates *S_1_* from *S_2_*. In the standard coalescent model, the probability of a mutation on a given branch is proportional to its length and independent of the coalescent process. Because branch lengths differ among gene trees, the probability of a DMI depends on the types of gene trees at a pair of loci. Conversely, the probability of a specific pair of gene trees at the two loci involved in a DMI (“DMI loci”) depends on the pattern of isolation. We can express the relationship between the probability of each pair of gene trees and the probability of an incompatibility through Bayes’ theorem.

Let *I* be the species pair(s) for which a DMI manifests – that is, *I* specifies the pattern of reproductive isolation between species for a DMI (Fig. 3). The probability that a pair of DMI loci have gene trees of type *T_x_* and *T_y_* respectively, can be expressed as

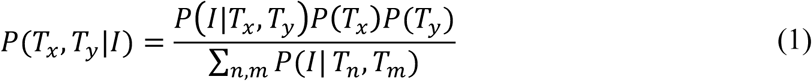
where *n* and *m* are each indices enumerating the four types of gene trees, and *P*(*T_x_*)*P*(*T_y_*) is the joint probability of *T_x_* and *T_y_* assuming independence as described above.

Assuming incompatibilities are rare between any given pair of loci, the conditional probability *P*(*I*\*T_x_*, *T_y_*) can be written as a sum of the probability that two mutations result in an incompatibility, across all branches of *T_x_* and *T_y_*. Let *x_α_* and *y_β_* be indexed branches on trees *T_x_* and *T_y_*, then

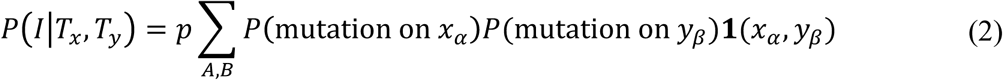
where **1**(*x_α_*, *y_β_*) is 1 when two mutations, on branches *x_α_* and *y_β_*, can generate a DMI with isolation pattern *I*, and 0 otherwise; and *p* is the probability of an incompatibility forming between untested allelic combinations (Orr 1995; Orr and Turelli 2001). This probability is valid when the mutations are independent, but the order in which mutations occur must be considered for derived-ancestral incompatibilities (see Methods and below). The probability of a mutation on a given branch can be estimated by its length,

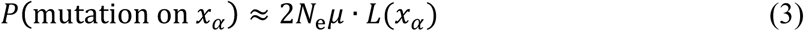
where 2*N*_e_*μ* is the population mutation parameter and *L*(*x_α_*) denotes the branch length of *x_α_* (Hudson 1992; see Supplemental Methods). The probability of a mutation on branch *y_β_* can similarly be estimated by its branch length.

With the probability of each pair of genealogies for a given pattern of reproductive isolation, we can calculate the probability of concordance with the species tree for a single DMI locus by summing the marginal probabilities for a concordant topology,

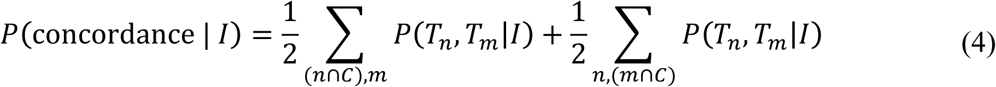
where *C* is the set of concordant gene tree indices. The probability of discordance can similarly be calculated from a sum of marginal probabilities.

#### Mutations that isolate sister taxa via two loci with discordant gene trees

Determining the limited branch segments on which incompatible alleles for a particular pattern of isolation can arise is central to the calculation of the conditional probability in Equation 2. In Figure 4, we illustrate the branch segments on which an incompatible allele isolating the sister taxa, *S_1_* × *S_2_*, can arise on two discordant gene trees, (*S_2_*, *S_3_*) *S_1_* and (*S_1_*, *S_3_*) *S_2_*. (For the segments on all gene tree pairs, see Appendix 1 of the Supplemental Materials.) We divide segments on the gene trees in Figure 4 by speciation and coalescent events, labeling each segment by its endpoints (e.g. *a-d* describes the segment specific to *S_1_* from the present to the most recent speciation event). Two mutations on the highlighted segments in each pair of gene trees in Figure 4, one on the left-hand tree and one on the right-hand tree, can produce alleles isolating *S_1_* and *S_2_*. For example, Figure 4a shows the potential for an *S_1_* × *S_2_* incompatibility from a derived allele that arises on the *a-d* segment of the left-hand tree (inherited by *S_1_*) and a derived allele that arises on segments *b-e, e-g*, or *g-k* (inherited by *S_2_*). Since we do not allow incompatibilities to arise before *S_1_* and *S_2_* diverge, one mutation must occur on a segment after divergence (i.e. a pair of incompatible alleles cannot arise, for instance, along segment *d-g* on the left-hand tree and *e-g* on the right-hand tree).

**Figure 4.**
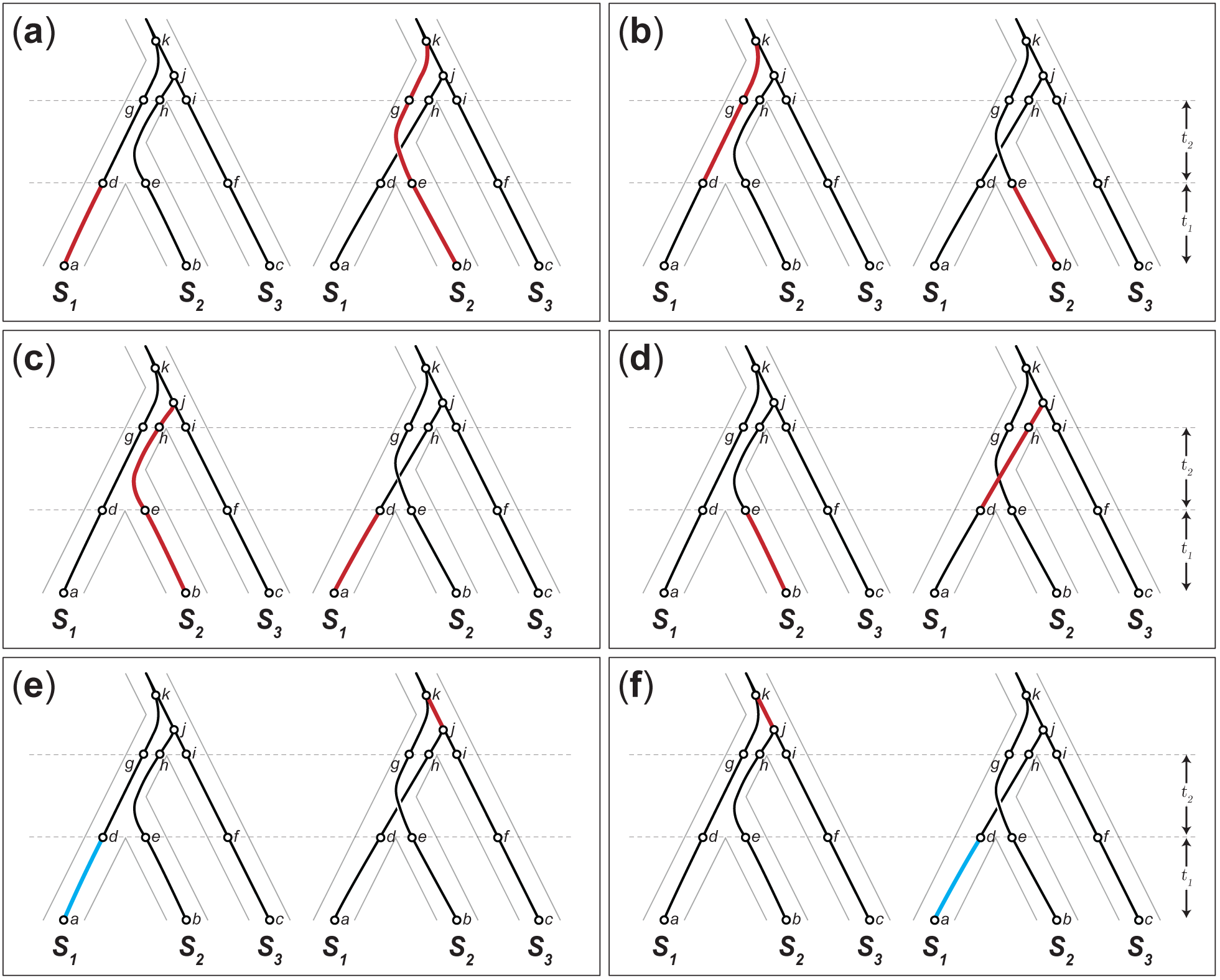
Branch segments on the discordant gene trees, (*S_2_*, *S_3_*) *S_1_* and (*S_1_, S_3_*) *S_2_*, that can give rise to incompatible alleles isolating *S_1_* and *S_2_*. In (a)-(d), a mutation on the left-hand tree, (*S_2_*, *S_3_*) *S_1_*, and a mutation on the right-hand tree (*S_1_*, *S_3_*) *S_2_*, can give rise to a derived-derived incompatibility. These panels show all combinations of branch segments on which mutations could give rise to a derived-derived incompatibility on the pair of discordant trees shown. Panels (e) and (f) show all combinations of branch segments on which mutations could give rise to a derived-ancestral incompatibility on this pair of trees. A mutation on branch segment *j-k* (red) leaves *S_2_* with an ancestral allele that can be incompatible with a derived allele produced by a mutation on branch segment *a-d* (blue) and inherited by *S_1_* (see main text).

Interestingly, loci with discordant gene trees can also produce derived-ancestral incompatibilities between sister taxa (Figs. 4e and 4f). In Figure 4e, a mutation on segment *j-k* on the right-hand tree produces a derived allele inherited by *S_1_* and *S_3_*. The second derived allele on segment *a-d* on the left-hand tree is inherited by *S_1_*, but arises in the background of the derived allele from the first mutation, creating the potential for an incompatibility with the ancestral allele on *S_2_*. A derived-ancestral incompatibility between sister taxa is only possible in a model that allows incompatible alleles to arise before divergence.

#### Discordance is more likely at loci isolating sister taxa

The probability of concordance with the species tree at a DMI locus in a three-species complex depends on the pattern of reproductive isolation considered. Figure 5 shows the probability of concordance for a DMI locus participating in each of the four possible patterns of isolation. The values depicted are relative to the expected probability of concordance for random, non-DMI loci: 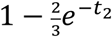 (Hudson 1983). The greatest deviations from this background probability of concordance occur when little time has elapsed since, and between, speciation events; this is true for all four patterns of isolation (Fig. 5). When *t_1_* and *t_2_* are short, branch segments on which incompatible alleles can arise vary greatly between the four types of gene trees (Fig. 1); this in turn leads to larger disparities in discordance among patterns of isolation (Fig. 5).

**Figure 5.**
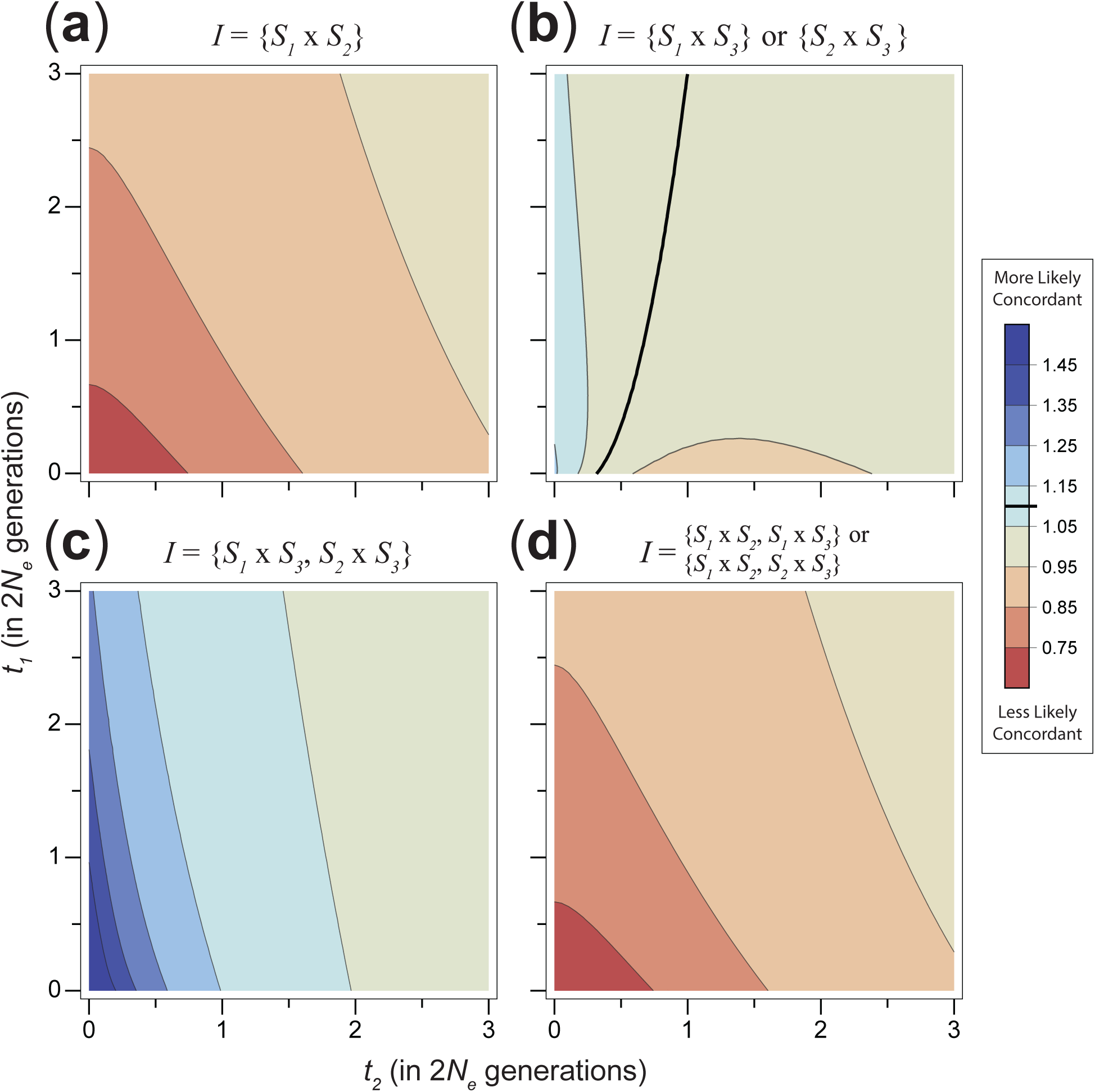
Relative probability of concordance conditioned on the pattern of reproductive isolation. Contour plots show the probability that a DMI locus has a concordant gene tree, relative to the expected probability of concordance for non-DMI loci. Bold line in (b) and in the legend indicate where the probability of concordance is equal to the expected value from a non-DMI locus. All other panels show results that are always either above or below a ratio of 1.

Two contrasting patterns emerge as time since divergence grows, depending on whether the locus participates in an incompatibility between sister taxa, *S_1_* × *S_2_*. For a locus participating in an incompatibility isolating sister taxa, the relative probability of concordance is at a minimum when *t_1_* and *t_2_* are short, and are 33% less likely to be concordant as these times approach zero (Fig. 5a, 5d). Loci that do not participate in an incompatibility between sister taxa are *more* likely to be concordant when times are short, up to 67% more likely to be concordant as times approach zero (Fig. 5b, 5c).

The contrast between isolation patterns derives from the restrictions placed on the position of mutations when conditioning on each pattern of reproductive isolation. On concordant trees, alleles involved in sister-taxa incompatibility must arise before (looking backward in time) the coalescence of lineages from the sister species. This coalescence is, on average, deeper on discordant trees, providing more time for the appropriate mutations to arise. Conversely, the deeper coalescence on discordant trees also reduces the shared branch length leading to sister species. The reduced potential for an incompatible allele shared between the sister species on discordant trees increases the chances that a shared incompatibility isolating both *S_1_* × *S_3_* and *S_2_* × *S_3_* is due to loci with concordant trees.

#### DMI loci are on average slightly more likely to be discordant

The results above were presented separately for the four different patterns of reproductive isolation among three species. In order to present the probability of discordance across all patterns of isolation, we must take into account the likelihood of each isolation pattern under different histories. For example, pairs of species that have been diverged longer are more likely to harbor incompatibilities, and thus, more likely to be among the species that are isolated. The likelihood that a DMI locus confers a specific pattern of isolation therefore depends on both tip length, *t_1_*, and internal branch length, *t_2_*. As a result, the general, unconditioned probability of concordance at a DMI locus depends on the likelihood of each isolation pattern.

We calculate the relative probability of each isolation pattern by conditioning on the observation of a DMI. The probability of a particular pattern of isolation, *I*_0_, can be written as

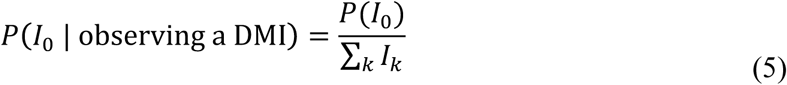
where *k* is an index for the patterns of isolation and *P*(*I_k_*) is the denominator in Equation 1,

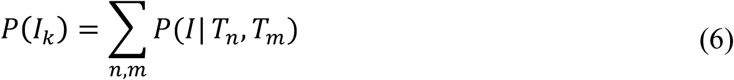

From this, we compare the relative probability of each isolation pattern in our model, which considers ILS, to a model on a fixed species tree with no ILS. For the model with a fixed species tree we use the expected number of incompatibilities from Wang et al. (2013) to compute the probability of each isolation pattern. In both models, isolation patterns that include an incompatibility between sister species are most likely when tip lengths, *t_1_*, are long relative to the inter-speciation time, *t_2_* (Figs. S2a, S2d; S3a, S3d). The opposite case, with short tip lengths relative to inter-speciation time, favors the isolation pattern where both sister species are incompatible with the third species (Figs. S2c, S3c). For intermediate values of *t_1_* and *t_2_*, the most likely case is an incompatibility that isolates one of the two more distantly related species pairs, that is, isolating *S_1_* × *S_3_* or *S_2_* × *S_3_* (Figs. S2b, S3b).

The introduction of ILS substantially increases the proportion of incompatibilities that isolate more than one species pair (i.e. isolation patterns (c) and (d) in Fig. 3). On a fixed species tree with no ILS, no more than one-third of incompatibilities ever isolate multiple species pairs, but in a model with ILS, more than half isolate multiple species pairs when *t_2_* is short relative to *t_1_* (Fig. S3). This difference arises from the additional branch length that is specific to one lineage on discordant topologies. Coalescence on discordant topologies can only occur in the ancestral population of all three species, substantially increasing lineage-specific branch lengths. When mutations that produce incompatible alleles are inherited by the same lineage, a derived-ancestral incompatibility forms with other lineages (Fig. S1). Discordant topologies increase the chances for these shared incompatibilities, especially when *t_2_* is short relative to *t_1_*.

Putting together the probability of each isolation pattern with its probability of concordance, the general, unconditioned probability of concordance can be calculated by the sum,

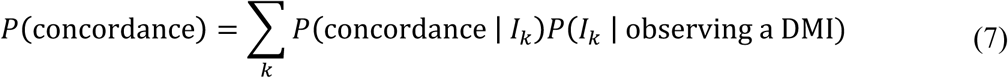

Figure 6a shows the general probability of species tree concordance for a DMI locus, across all patterns of isolation. When tip lengths, *t_1_*, are short, concordance is slightly more likely, up to 27% as both times approach 0. However, for most combinations of times, *t_1_* and *t_2_*, DMI loci are slightly less likely to be concordant. Overall, gene trees for a locus participating in a DMI have a probability of concordance very slightly below the background value of 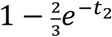.

**Figure 6.**
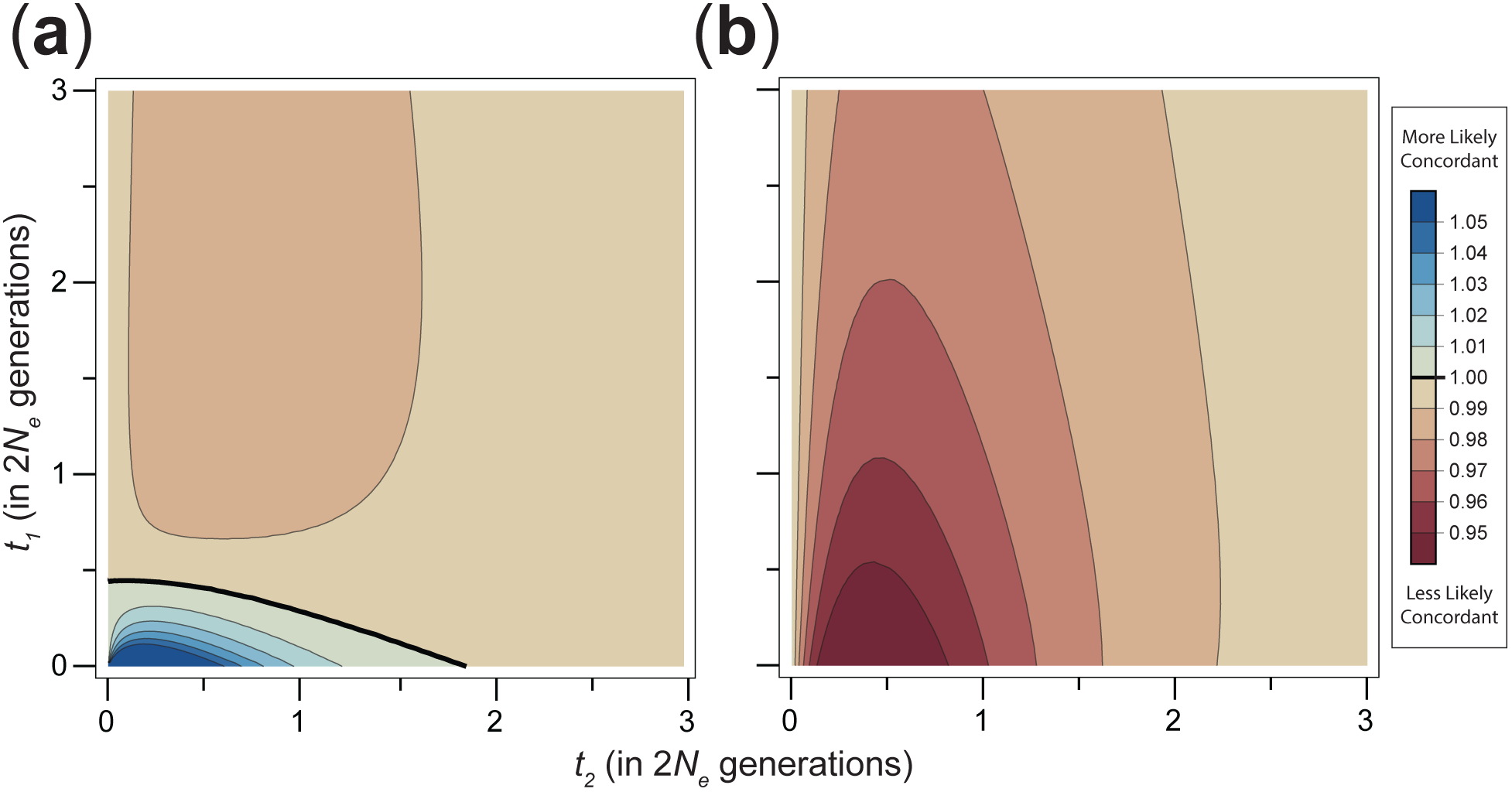
Relative probability of concordance for a DMI locus. (a) Probability of concordance for a DMI locus in a model restricting incompatibilities from arising in the same population. Concordance is slightly more likely when times are short. Bolded line shows the contour where concordance is equal to the canonical expectation from coalescent theory. (b) Probability of concordance for a DMI locus in a model allowing incompatibilities to arise in the same population. Concordance is always less likely.

#### Allowing incompatibilities to arise in the same population

Our model of DMI loci featuring discordant gene trees has, thus far, explicitly prevented incompatibilities from arising in the same population. This prohibition assumes that selection against incompatibilities is strong enough to prevent the persistence of incompatible alleles in a population. Evidence for the variability of reproductive isolation within populations suggests that the strength of selection may be insufficient to prevent polymorphic incompatibilities from existing (e.g. Corbett-Detig et al. 2013). To address this possibility in our model, we relax this prohibition, allowing incompatible alleles to arise and segregate in ancestral populations as long as extant lineages do not individually carry the incompatible genotype.

Because our model considers the genealogical history of a DMI locus, incompatibilities that arise in the same population can be incorporated with relative ease. The restriction against these incompatibilities has been enforced by requiring at least one derived allele in an incompatibility to arise after divergence between species pairs. This restriction can be lifted by allowing derived alleles to arise up to (backward in time) the point of coalescence. For derived-ancestral incompatibilities, we continue to enforce the restrictions from mutation order. That is, a derived allele participating in a derived-ancestral incompatibility may only arise after a mutation has already produced the compatible allele (see Supplemental Methods). The probability of concordance when considering incompatibilities that arise within populations before divergence can then be calculated by using a relaxed indicator function in Equation 2. (The indicator function of each gene tree pair is available in Appendix 2 of the Supplemental Materials.)

Overall, allowing the unrestricted emergence of incompatibilities within populations reduces the probability that DMI loci will have concordant gene trees (Fig. 5b). When tip lengths, *t*_1_, are short, the probability of concordance can be reduced by up to 7% relative to the background expectation. In contrast to the model that restricts DMIs from emerging before divergence, concordance is always less likely than the expectation for non-DMI loci.

A model that allows incompatibilities to arise within populations also increases the probability of DMIs that isolate sister species (Figs. S4a, S4d). This results from the additional opportunities for isolating mutations on inner branch segments. Unlike incompatibilities isolating more distantly related species pairs, incompatible alleles on inner branch segments were consistently restricted among sister species from producing incompatibilities within populations. This model also causes the pattern of reproductive isolation to have an even greater impact on the probability of concordance. Qualitatively, the patterns of concordance conditioned on each isolation pattern are similar (see Fig. S5). However, loci participating in incompatibilities between sister taxa, *S_1_* × *S_2_*, are much less likely to be concordant, up to 66% less likely as *t_1_* and *t_2_* approach 0. Meanwhile, a locus participating in an incompatibility shared between species pairs *S_1_* × *S_3_* and *S_2_* × *S_3_* is up to 133% more likely to be concordant. As before, the differences in branch length and topology between the types of gene trees are greatest when *t_1_* and *t_2_* are short, but these differences are amplified when incompatibilities can arise in the same population before divergence.

#### Incompatible alleles are likely to have arisen in ancestral populations

In the previous section we allowed pairs of incompatible alleles to arise in the same population before divergence. Because of selection against incompatibilities within populations, such a history for DMI loci should be less common. However, an incompatible allele in a DMI pair could have arisen in an ancestral population without ever having caused an incompatibility within populations. Such an allele would have arisen in the ancestral population and then fixed in one lineage before becoming incompatible with a new mutation after divergence (e.g. the pair of DMI loci in Fig. 4d). As taxa spend more time diverged, incompatibilities between them become more likely to be the result of interactions between new mutations that arise post-divergence. In contrast to this scenario, many formulations of the DM model only allow incompatibilities to form from new mutations after divergence (Orr 1995; Orr and Turelli 2001; Fierst and Hansen 2010; Livingstone et al. 2012; Wang et al. 2013; Fraisse et al. 2014).

To examine the extent to which an incompatible allele is likely to have arisen prior to speciation, we consider its probability in our DM model with ILS. Mutations that occur before divergence give rise to incompatible alleles (but not incompatibilities) in ancestral populations. Branch segments on gene trees positioned before divergence bear such mutations. We can calculate the probability that an incompatibility involves ancestrally arising alleles by adjusting Equation 2 to count only mutations on pre-divergence branch segments,

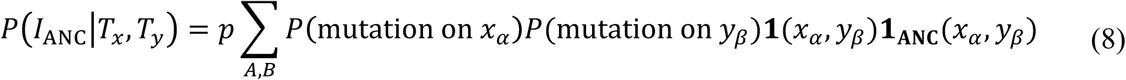
where **1_ANC_**(*x_α_*, *y_β_*) is 1 when either branch segment is positioned before divergence and 0 otherwise (see Supplemental Materials). The unconditional probability that an incompatibility involves one ancestrally arising allele can then be calculated following Equation 5. Figure 7 shows the proportion of incompatibilities in a three-species complex involving one incompatible allele that arose in an ancestral population. As branch lengths increase, incompatibilities are more likely to form solely from alleles that arose after divergence.

**Figure 7.**
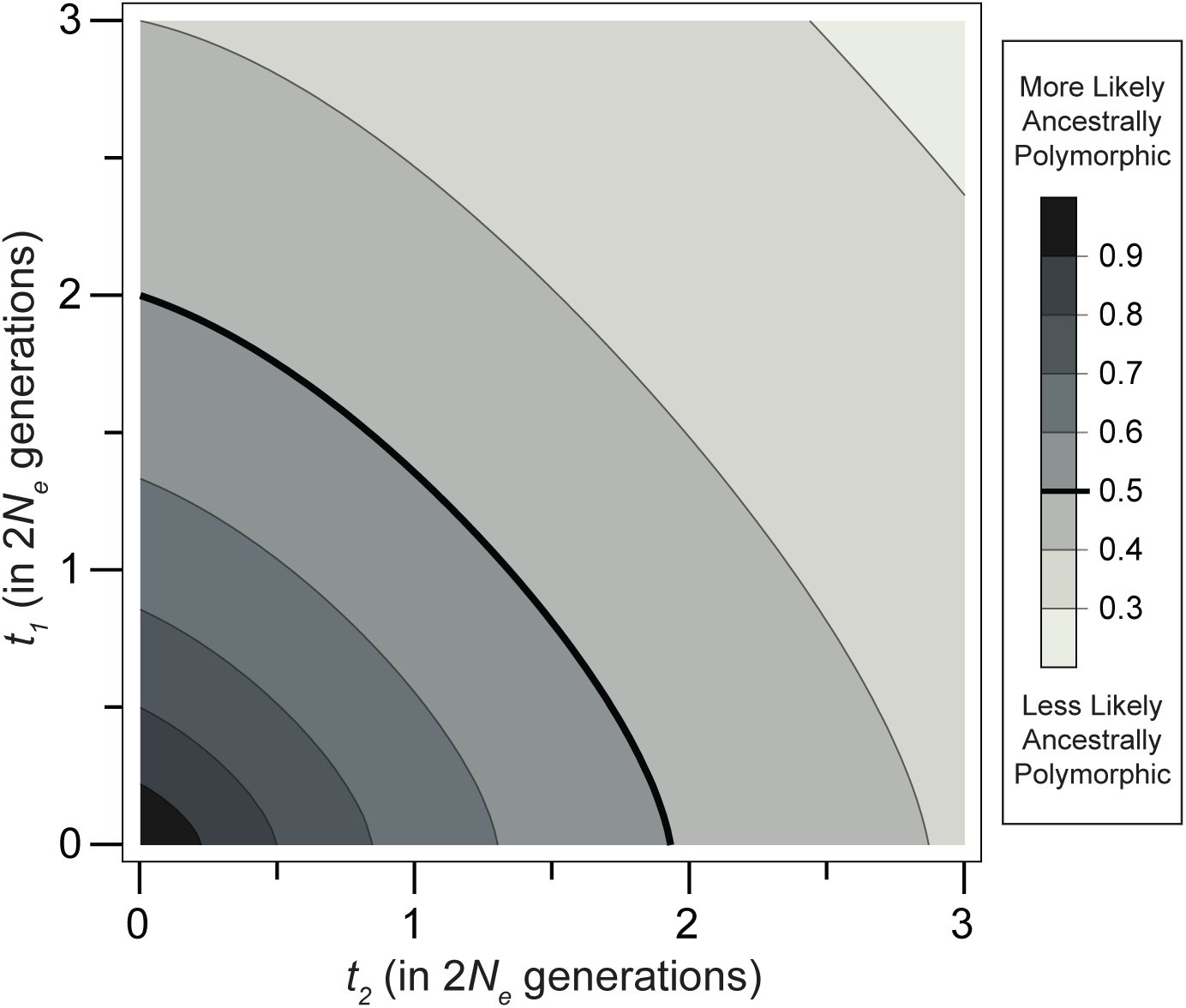
Probability that an incompatibility involves an allele that arose prior to speciation. Bold line shows the contour where a pairwise DMI is equally likely to form from at least one ancestrally arising mutation as from alleles appearing completely after divergence.

The pattern of isolation at a DMI has a substantial influence on whether ancestrally arising alleles participate in the incompatibility (Fig. S6). When considering incompatibilities between sister taxa, *S_1_* × *S_2_*, the proportion of incompatibilities that have at least one ancestrally arising allele depends only on tip length, *t_1_*, and effective population size, *N*_e_. This is because the branch length on which incompatible alleles can arise in the ancestral population depends only on the expected time to coalescence of two lineages from the point of their divergence, which equals 2*N*_e_. The probability of an ancestral allele in a DMI for this case is proportional to the product of the two pre-divergence segments unique to each lineage and the post-divergence segment of its counterpart (see Fig. S7); this is equal to 2 · 2*N*_e_ · *t_1_*. Meanwhile, the probability that a DMI involves only mutations arising post-speciation is proportional to the product of the post-divergence lengths, *t_1_*^2^. For other patterns of isolation, the probability that ancestral alleles participate in the DMI decreases with inter-speciation time, *t_2_*.

Overall, for examined speciation times, incompatibilities are likely to involve at least one ancestrally arising allele. When incompatibility loci are allowed to arise in the same population, the probability that incompatibilities result from ancestrally arising alleles increases, but the qualitative patterns are not substantially different from those described above (Fig. S8).

## Discussion

Our results show that DMI loci are slightly more likely to be discordant because discordant trees offer more opportunities for the formation of incompatible alleles. This finding follows inevitably from the fact that the mutational target size for a DMI at a particular locus depends on its gene tree, coupled with the constraint that incompatible alleles in a pairwise DMI can only arise on certain pairs of branch segments. A DMI locus is more likely to have a discordant genealogy if these branch segments are longer on discordant trees, and vice versa. The relevant branch segments differ for different patterns of isolation, such that alleles isolating sister taxa are more likely to form on discordant trees, while alleles isolating both sister taxa from a third taxon are more likely to form on concordant trees. Our results are in opposition to previous verbal models (e.g. Ting et al. (2000)), and suggest that gene tree concordance is unlikely to be useful for identifying DMI loci across the genome. Given the slight excess of discordance expected at DMI loci, gene tree discordance is also unlikely to be useful for this task.

Our findings are specific to a neutral model with only ILS acting, but other factors could plausibly cause speciation loci to be more informative about species relationships. There are two further arguments used to support the claim that speciation loci are more likely to have gene trees that are concordant with the species tree. The first argument concerns the potential for gene flow between incipient species. At loci conferring reproductive isolation, gene flow should be reduced by the lower fitness of hybrids that inherit the incompatible genotype. While substantial gene flow between two species may scramble the genealogical history for the rest of the genome, DMI loci should retain their original history. There is no guarantee, however, that the original history of the locus is one that is concordant with the species tree. This is especially true as the number of species in the topology increases, precisely the case for which concordance would be relevant. DMI loci may indeed be more likely to be concordant than introgressed loci, but not any more so than other non-introgressed loci. Furthermore, DMI loci that do not provide a selective advantage in the lineage on which they arose are unlikely to persist in the presence of gene flow (Gavrilets 1997; Bank et al. 2012).

The second argument relies on speciation loci being the target of positive selection. Loci involved in DMI often bear the signature of positive selection (Coyne and Orr 2004; Orr et al. 2006). The interpretation is that incompatible alleles form from the rapid divergence of adaptively evolving loci. A history of positive selection at a locus increases the chances that it will have a concordant gene tree because selective sweeps reduce *N*_e_, and consequently reduce the time to coalescence (Kaplan et al. 1989). The frequency with which DMI loci are more likely to be concordant then depends on how often they are targets of positive selection. However, the higher chance of concordance from positive selection is not unique to loci that confer reproductive isolation; any locus that has experienced linked selection is more likely to be concordant with the species tree (Slatkin and Pollack 2006; Stukenbrock et al. 2011; Duthiel et al. 2015). An examination of loci likely to be affected by linked selection is a more direct way of gaining this insight into species relationships (e.g. Scally et al. 2012; Pease and Hahn 2013; Munch et al. 2016).

The model introduced in this manuscript for hybrid incompatibilities incorporates the coalescent and ILS, inheriting several limitations and assumptions. New mutations are assumed to be neutral and modeled as arising at a fixed rate from populations that have a stable effective population size. As in the traditional DM model, incompatibilities are assumed to have an equal probability of forming between untested allelic combinations, which arise at independent, unlinked loci. An important simplification for our model is the consideration of only a single history for an incompatibility locus in each lineage; that is, we consider only a haploid history for incompatibility loci. Among diploids (and systems with higher ploidy), this simplification is equivalent to assuming that incompatible alleles have fixed in their extant lineages without having passed through the incompatible genotype. Though incompatibility loci can be polymorphic in both extant and ancestral populations (Cutter 2012), tracking this polymorphism in extant populations would require the consideration of multiple lineages in each species. This would necessitate a model of dominance and fitness for each genotypic combination of incompatibility loci. Such a model is outside the scope of our focus here on ILS and the stochastic accumulation of incompatible alleles, but could be interesting for future work.

In examining the possible histories for incompatible alleles, our results highlight how likely they are to have arisen prior to speciation (i.e. in ancestral populations) when they do not affect fitness in conspecific backgrounds. Our results suggest that pairwise DMIs are likely to involve at least one incompatible allele that arose ancestrally until 3–4 *N*_e_ generations after species divergence. That incompatible alleles are more likely to be from post-speciation mutations as species diverge may not be surprising from a population genetics perspective, but this result may help to clarify an argument on the relative importance of derived versus ancestral alleles in incompatibilities (Cutter 2012). Derived alleles are expected to play a larger role in the formation of incompatibilities because each pairwise incompatibility must involve at least one derived allele (Orr 1995). The distinction between derived and ancestral alleles should, however, not be confused with the genealogical history of participating loci. Mutations that yield incompatible derived alleles can arise both before and after populations diverge; that is, “ancestral” alleles are not derived alleles that arose in ancestral populations. Similarly, a derived-ancestral incompatibility can be the product of two mutations along the same lineage after divergence. Whether an incompatible allele arose in the ancestral population of two or more species depends on the timing of the mutation rather than its state relative to a common ancestor. Thus, the percentage of incompatibilities that involve alleles that arose in ancestral populations decreases with time, but the percentage of ancestral alleles participating in incompatibilities (from derived-ancestral interactions) remains the same.

The identification of loci participating in hybrid incompatibilities between multiple closely related species pairs has spurred many new analyses, including phylogenetic comparisons among species (Cattani and Presgraves 2009; Scarpino et al. 2013; Sherman et al. 2014; Wang et al. 2015). These comparative approaches can help to elucidate the timing and progression of reproductive isolation (Moyle and Payseur 2009). But the analysis of incompatibilities between multiple species pairs also introduces genealogical ambiguity at incompatibility loci. When incompatible alleles in a DMI arise on gene trees with different topologies, identifying which branch of the species tree they arose on becomes challenging. In fact, incompatible alleles arising on discordant trees may be mapped onto the wrong branches of the species tree by standard methods. While a derived allele in a pairwise DMI will always be inherited by at least one of the lineages isolated by the incompatibility, the other allele can originate on branches that are shared with uninvolved lineages. For example, in the absence of ILS an incompatibility isolating only the sister species among three taxa (Fig. 3a) would be interpreted as the result of two mutations, each uniquely inherited by one of the sister species. However, in the presence of ILS such an incompatibility could involve a mutation inherited by one of the sister species and a third taxon (as in Fig. 4c, 4d). Discordance between gene trees is most pronounced when little time separates speciation events, and this is often the case for model systems of speciation, where the ability to perform interspecific crosses is a useful tool for genetic investigation (e.g. True et al. 1996; Slotman et al. 2004; Sweigart et al. 2006; Moyle and Nakazato 2008; Matute and Coyne 2010; White et al. 2011). Under these circumstances, inferences about the origin of incompatibilities from comparative mapping need to be mindful of the genealogical history at DMI loci.

## Materials and Methods

### Calculating branch lengths

To calculate the probability of concordance at a DMI locus, we find the probability of concordance conditioned on a particular pattern of isolation, *I*. This requires calculating the branch lengths from each pair of gene trees, and the indicator function representing the opportunity to produce an incompatibility (Eq. 2). The indicator function can be represented as a matrix, **1***_Tx,Ty_*, with rows and columns corresponding to the branches of the respective gene trees *T_x_* and *T_y_*. With four types of gene trees, there are ten unique matrices (see Appendix 1; Supplemental Material).

Branch segments on each of the four types of gene trees are labeled in Figure 8. The probability of a mutation on the branch segments of each tree, *T*, can be assembled into the vector *B_T_*. We order the lengths of each segment (using the segment names as labels for their expected lengths for convenience, i.e. *ij* = *L*(*i-j*)) in the vectors as follows,

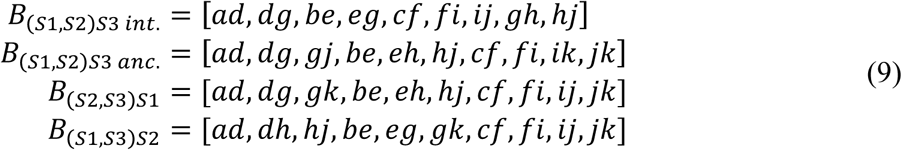

**Figure 8.**
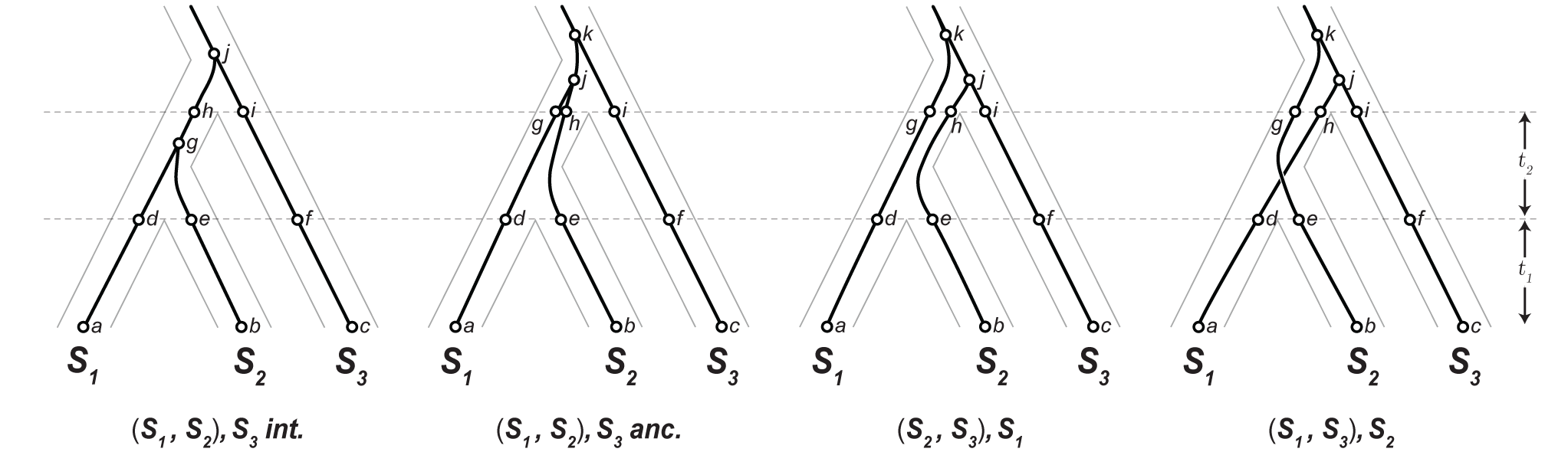
Four types of gene trees with branch segments labeled. Concordant trees are divided into those that coalesce in the time between species divergences (**inter**-) and those that coalescence in the **anc**estral population.

The length of segments whose ends are not coalescent events are simply *t_1_* or *t_2_*. Segments that coalesce in the ancestral population of all three species have expected lengths (in 2*N*_e_ coalescent units) of 1/3, 1, and 4/3, corresponding to the time to coalescence from three-to-two, two-to-one, and three-to-one lineages respectively.

Several segments on the (*S_1_, S_2_*) *S_3_ int*. tree have expected values that are conditioned on the first coalescence occurring by *t_2_*. This condition distinguishes this type of tree from the ancestrally coalescing tree with the same topology. Let *v* be a random variable from 0 to *t_2_*, representing the time from the most recent population divergence to the first coalescence event in the (*S_1_, S_2_*) *S_3_ int*. tree. The probability distribution function of *v* is given by the exponential distribution function for the coalescence of two lineages, *e*^−^*^t^*, divided by the probability of coalescence by *t_2_* (see Mendes and Hahn (2017)). Thus,

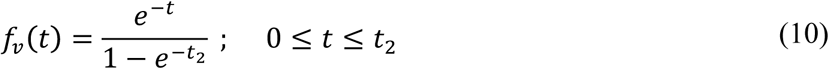

Let *q*(*t_2_*) be the expected value of *v* for a given value of *t_2_*, then the expected time from the first population divergence to the first coalescence in the (*S_1_, S_2_*) *S_3_ int*. tree is

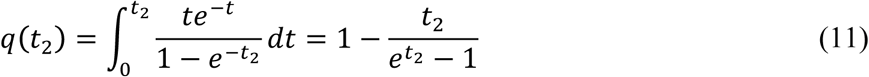

Filling in the segment lengths from Eq. 9, the values for *B_T_* become

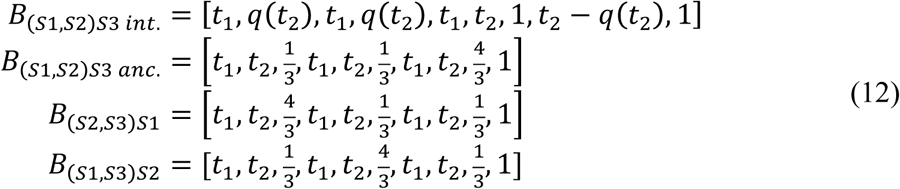

With the terms in Eq. 2 written as vectors and matrices, *P*(*I*\*T_x_*, *T_y_*) can be written conveniently as the matrix product

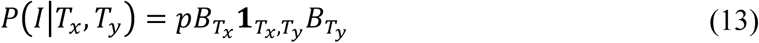

For four gene trees, there are 16 combinations of *T_x_, T_y_*. We form a 4x4 matrix, ***D***(*I*), for each isolation pattern *I*, whose entries are the above matrix product (Eq. 13) for each gene tree pair. The rows and columns in ***D***(*I*) are ordered so that they are consistent with the order of trees in Figure 8.

To arrive at the probability of each gene tree pair, Bayes’ theorem is applied (Eq. 1). The numerator from Eq. 1 can be calculated from each element of the ***D***(*I*) product matrix, multiplied by the unconditional probability of each gene tree pair. The probability for each type of gene tree (Fig. 1) can be assembled into a vector of tree probabilities, *T_prob_*, as

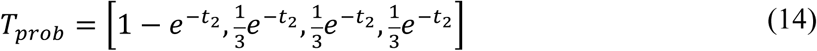

The numerator in Eq. 1 for each gene tree pair can then be represented as the element-wise product of ***D***(*I*) with the outer product of *T_prob_* with itself. We call this 4x4 matrix ***P***(*I*),

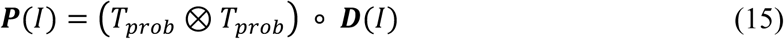

The denominator for Eq. 1 is the sum of all elements in the ***P***(*I*) matrix.

Returning to Eq. 4, the probability of concordance for a single DMI locus is given by the sum of all elements in the first two rows and columns of **P**(*I*), divided by twice the sum of all elements. Note that the parameters *p* and *2N*_e_*_μ_* do not appear in the probability of concordance as they are cancelled by the denominator in Eq. 1.

#### Mutation order and derived-ancestral incompatibilities

Thus far, the probability of each mutation in a DMI pair has been treated as an independent event. That is, the joint probability is calculated as a product of the mutation probabilities on each branch (Eq. 2). While this assumption is valid for derived-derived incompatibilities, greater care must be taken for derived-ancestral incompatibilities due to the order in which mutations must occur. Consider a potential derived-ancestral incompatibility between a mutation on segment *g-h* of the (*S_1_, S_2_*) *S_3_ int*. tree and a mutation on segment *d-g* of the (*S_2_*, *S_3_*) *S_1_* tree, as in Figure 9.

**Figure 9.**
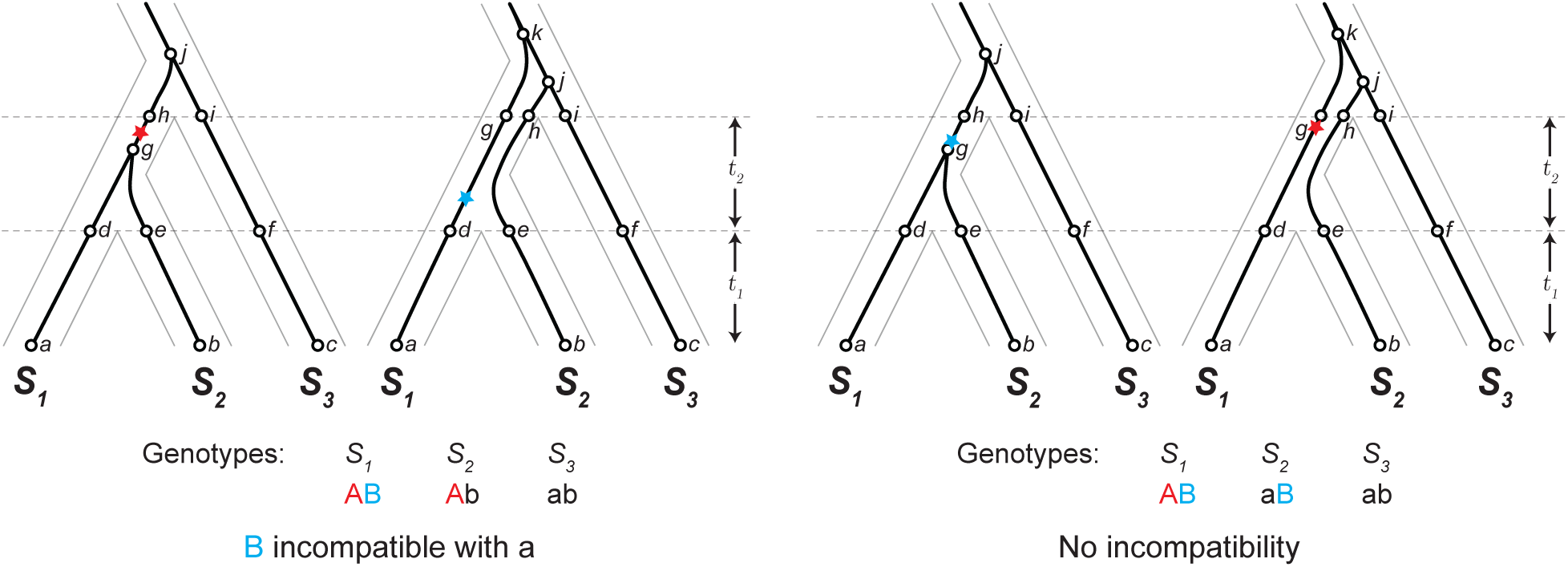
Mutation order determines the potential for a derived-ancestral incompatibility. Red and blue stars mark the position of the first and second mutations producing incompatible alleles. The ancestral genotype for the two loci is denoted ‘ab’, with the mutations producing derived alleles ‘A’ and ‘B’. (a) Derived-ancestral incompatibility from the derived allele in the *S_1_* lineage with the ancestral allele in the *S_3_* lineage. (b) No incompatibility forms because the derived-ancestral genotype persists in the *S_2_* lineage.

A mutation on segment *g-h*, followed by a mutation on *d-g* can result in a derived-ancestral incompatibility between *S_1_* and *S_3_*. The converse, a mutation that occurs first on segment *d-g* followed by a mutation on *g-h*, cannot result in a derived-ancestral incompatibility because the ancestral allele persists in the *S_2_* lineage. Since the incompatible allelic combination already exists in *S_2_*, this combination cannot be the cause of an incompatibility between *S_1_* and *S_3_*.

This asymmetry from mutation order occurs because the derived allele in a derived-ancestral incompatibility must arise from the second mutation. If the derived allele arose from the first mutation, it would immediately produce the incompatible genotype. In the previous example, when the second mutation arises on segment *d-g*, the derived allele is inherited by the *S_1_* lineage; whereas when the second mutation arises on segment *g-h*, both the *S_1_* and *S_2_* lineage inherit the derived allele. Between branch segments that have contemporaneous endpoints, this asymmetry does not exist and the order of mutations does not matter. For example, the first mutation on segment *a-d* of either tree in Figure 9 leads to an ancestral allele that can be incompatible with a second mutation at segment *a-d* on the other tree.

Differences in mutation order produce this asymmetry only when the inheritance of the derived allele is made ambiguous by the timing of a coalescent event. This is only possible when the mutations are from segments that overlap temporally. Most pairs of segments where mutation order is ambiguous occur before any population divergence has occurred (Fig. 8). When incompatibilities are restricted to arising only after populations diverge, these segments do not cause any incompatibilities. The only gene tree pairs where mutation order must be considered in this model are those that involve the (*S_1_*, *S_2_*) *S_3_ int*. tree. Here, the coalescence in the ancestral population of *S_1_* and *S_2_* can change the identity of the derived allele in an *S_1_* × *S_3_* incompatibility and an *S_2_* × *S_3_* incompatibility.

Returning to our earlier example, we calculate the joint probability of two mutations leading to a derived-ancestral incompatibility from segments *g-h* on an (*S_1_*, *S_2_*) *S_3_ int*. tree and *d-g* on an (*S_2_*, *S_3_*) *S_1_* tree. Such an incompatibility requires the first mutation to be on segment *g-h*, restricting the timing of the second mutation on *d-g*. The probability of the first mutation is unrestricted and is equal to the branch length of *g-h*, which has an expected value of *t_2_* − *q*(*t_2_*). Let *τ* be the time from the *S_1_, S_2_* divergence to the first mutation on segment *g-h*. Assuming that a single mutation occurs on segment *g-h*, the probability of the first mutation should be uniformly distributed along the length of segment *g-h*. Then,

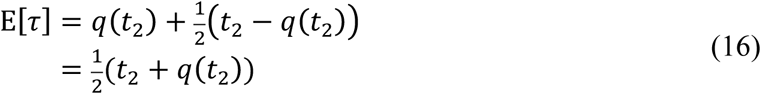

Mutations that occur before the first mutation on *g-h* do not cause an incompatibility, thus the expected value of *τ* is also the expected length of the sub-segment of *d-g* from which a second mutation may arise. The probability of an incompatibility from these two segments can then be calculated from the product of E[*τ*] and *t_2_* − *q*(*t_2_*).

This probability applies for each of the gene tree pairs involving (*S_1_*, *S_2_*) *S_3_ int*. and any other gene tree for producing a derived-ancestral incompatibility between *S_1_* and *S_3_*. Similarly, the same reasoning can be applied for interactions between mutations on *g-h* and *e-h* for incompatibilities between *S_2_* and *S_3_*. When both gene trees are (*S_1_*, *S_2_*) *S_3_ int*., the calculation is more involved due to the coalescence at the end of branch segments from both trees (see Supplemental Material).

## Supplemental Material

### Supplemental Figures

**Figure S1.**
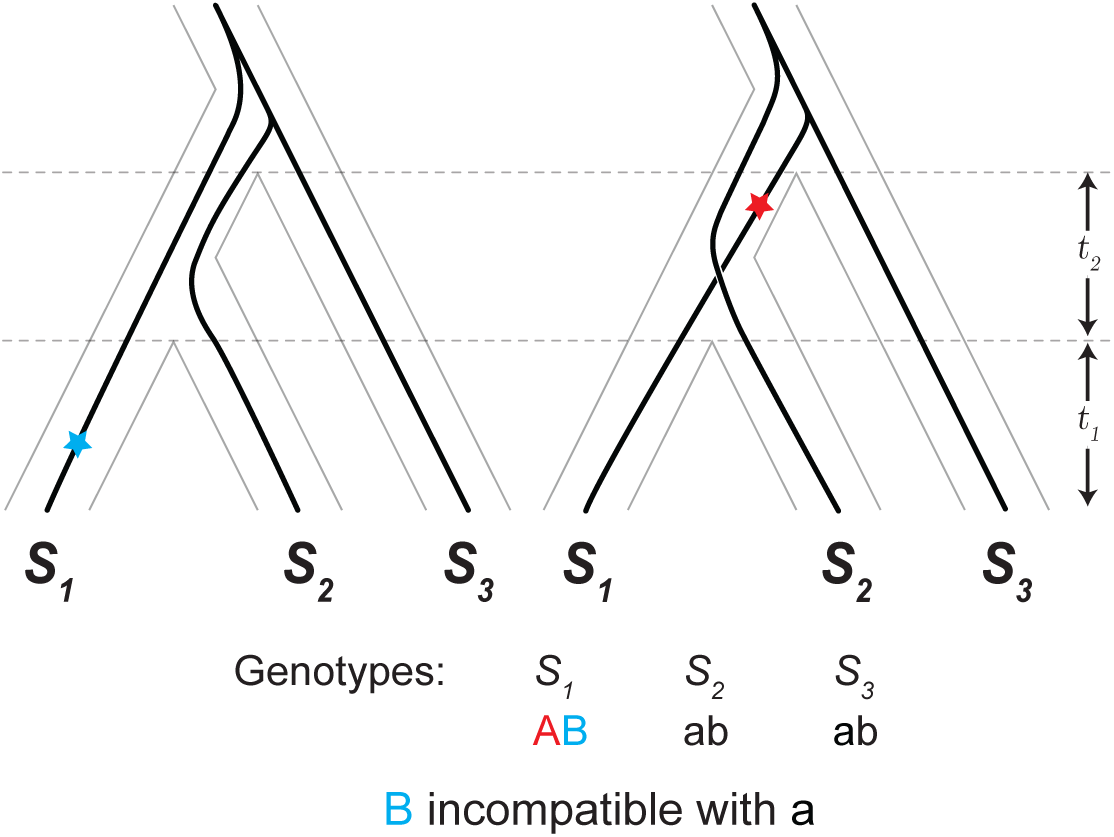
Derived-ancestral incompatibility shared between *S1* × *S2* and *S1* × *S3* due to a shared ancestral allele. Red and blue stars mark the position of the first and second mutations, respectively, that produce incompatible alleles. The ancestral genotype, ‘ab’ is present in lineages *S2* and *S3*, and incompatible with the ‘AB’ genotype in lineage *S1*.

**Figure S2.**
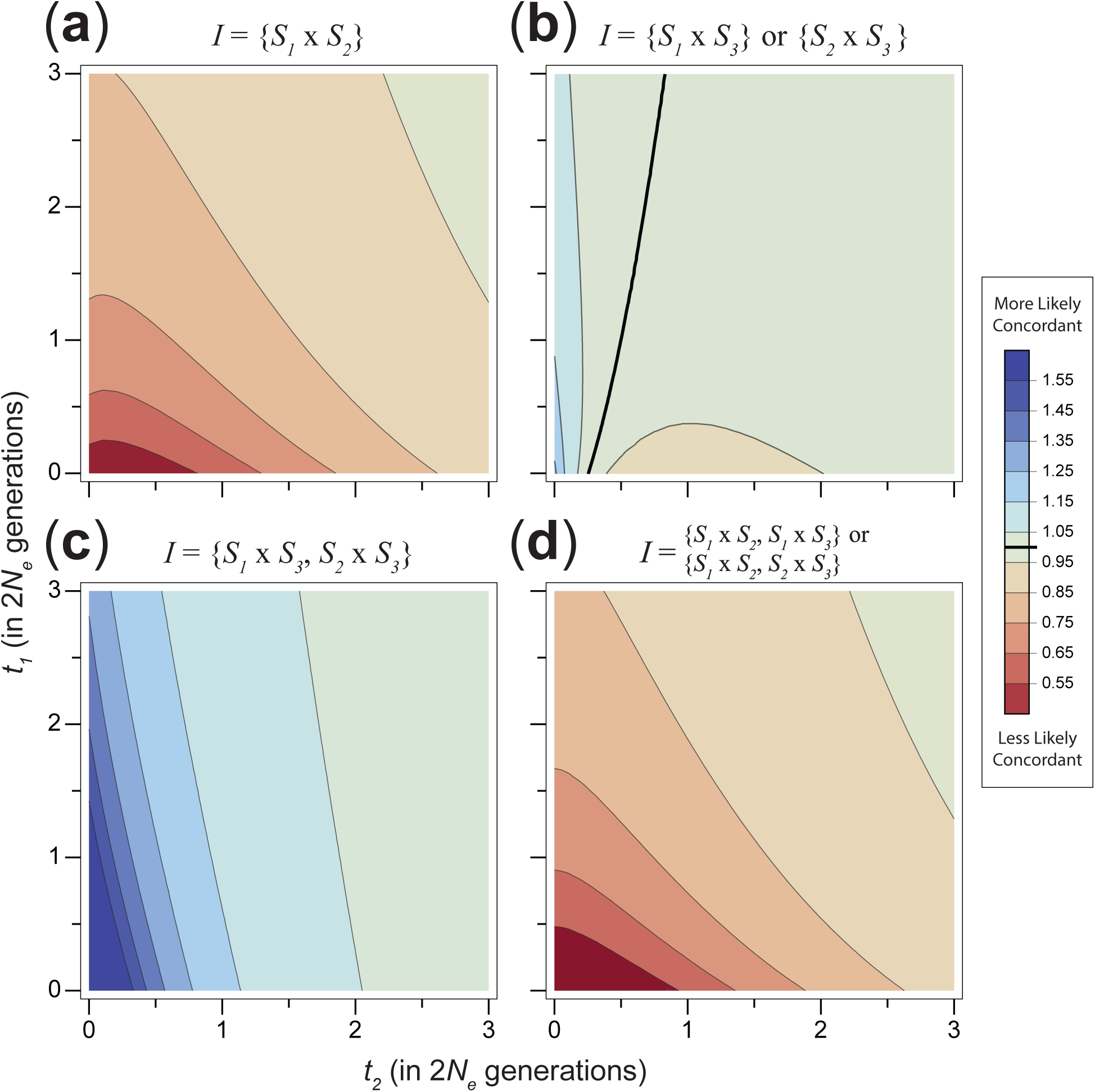
Relative probability of concordance conditioned on the pattern of reproductive isolation (polymorphic incompatibilities allowed). Bold line in (b), and in the legend, indicates where the probability of concordance is equal to the expected value from a non-DMI locus. All other panels show results that are all either above or below a ratio of 1.

**Figure S3.**
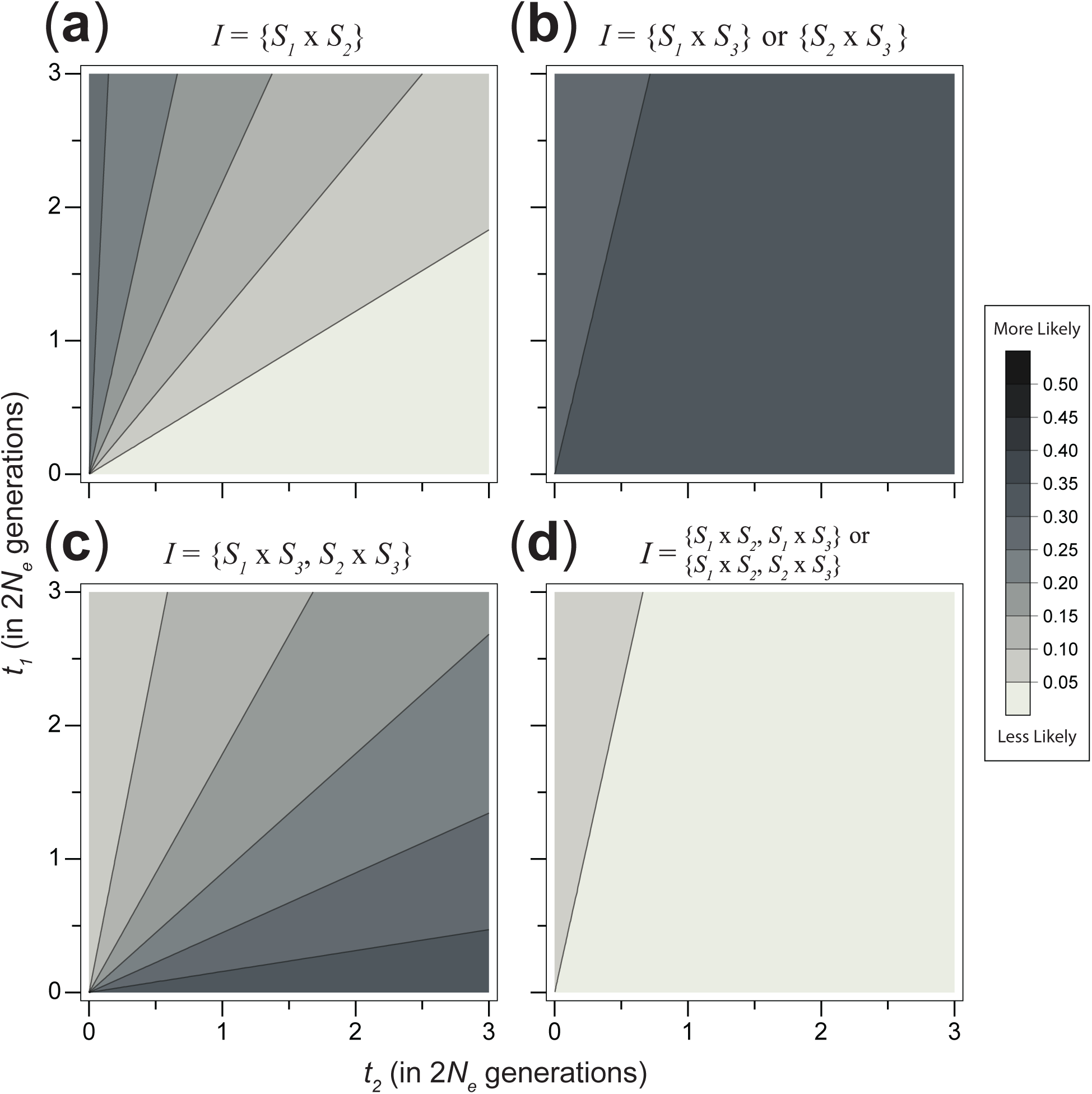
Probability for different patterns of reproductive isolation; model of DMI with loci on a fixed species tree.

**Figure S4.**
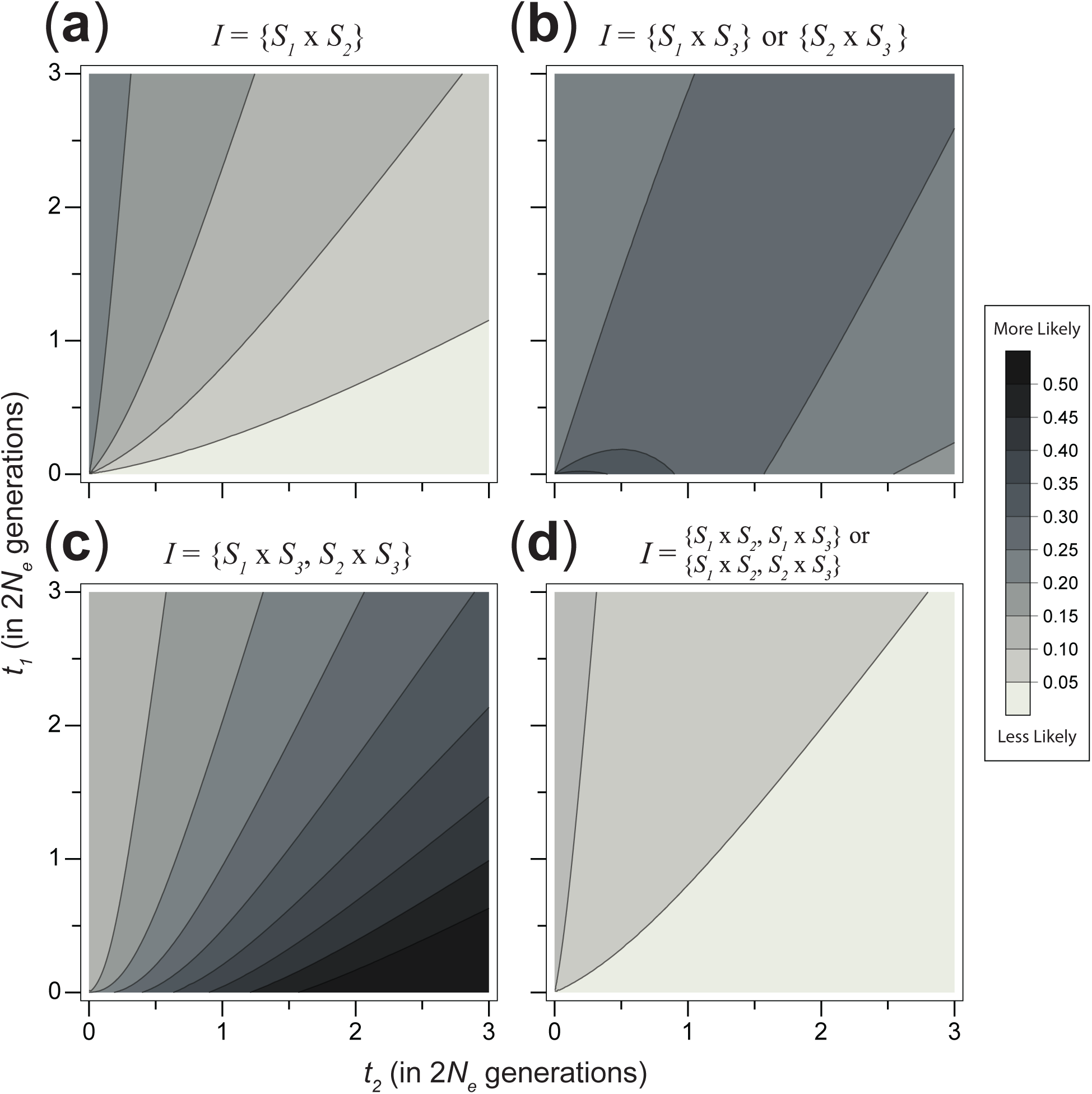
Probability for different patterns of reproductive isolation; model of DMI with loci subject to ILS.

**Figure S5.**
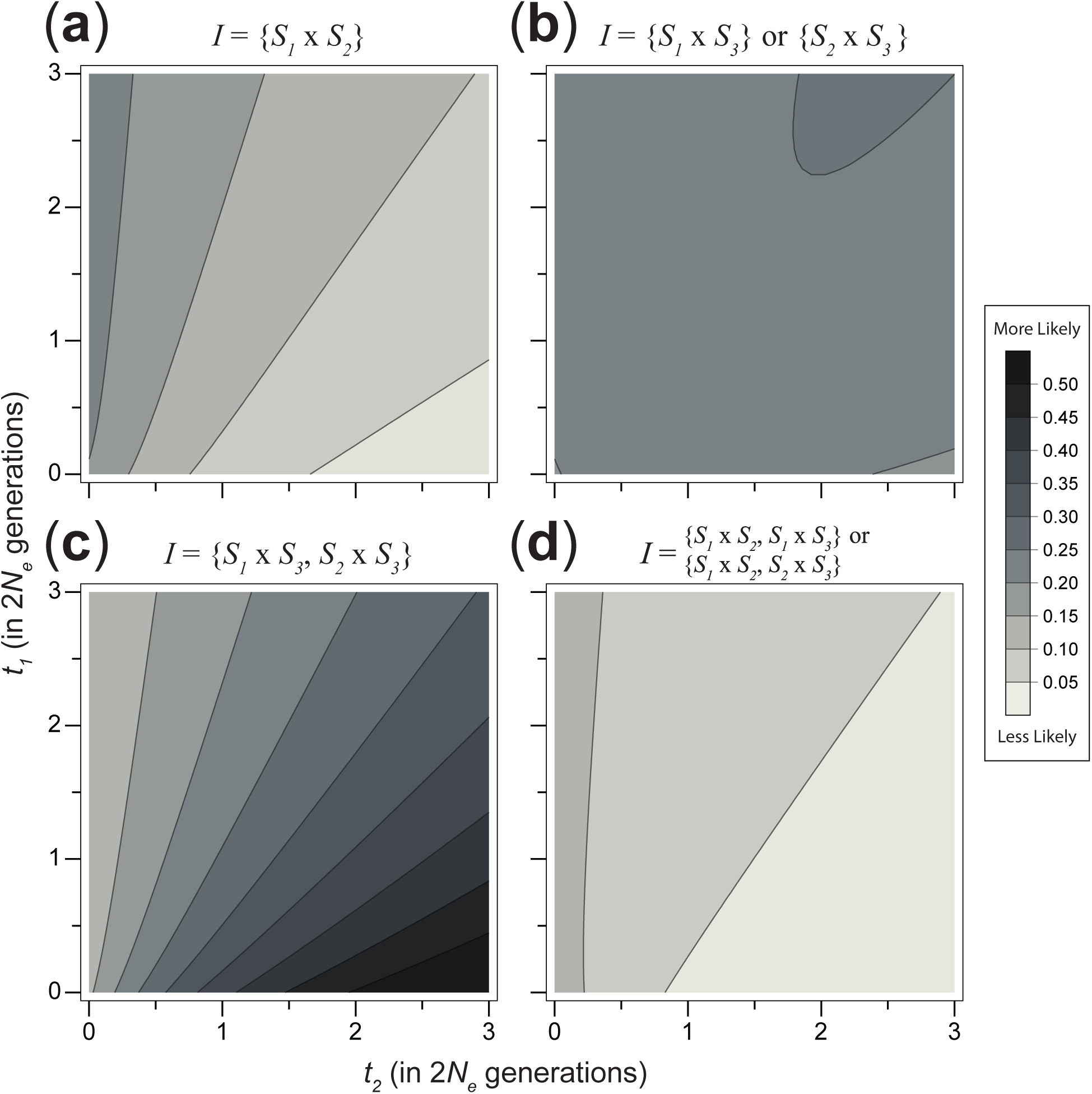
Probability for different patterns of reproductive isolation; model of DMI with potentially polymorphic loci, in addition to being subject to ILS.

**Figure S6.**
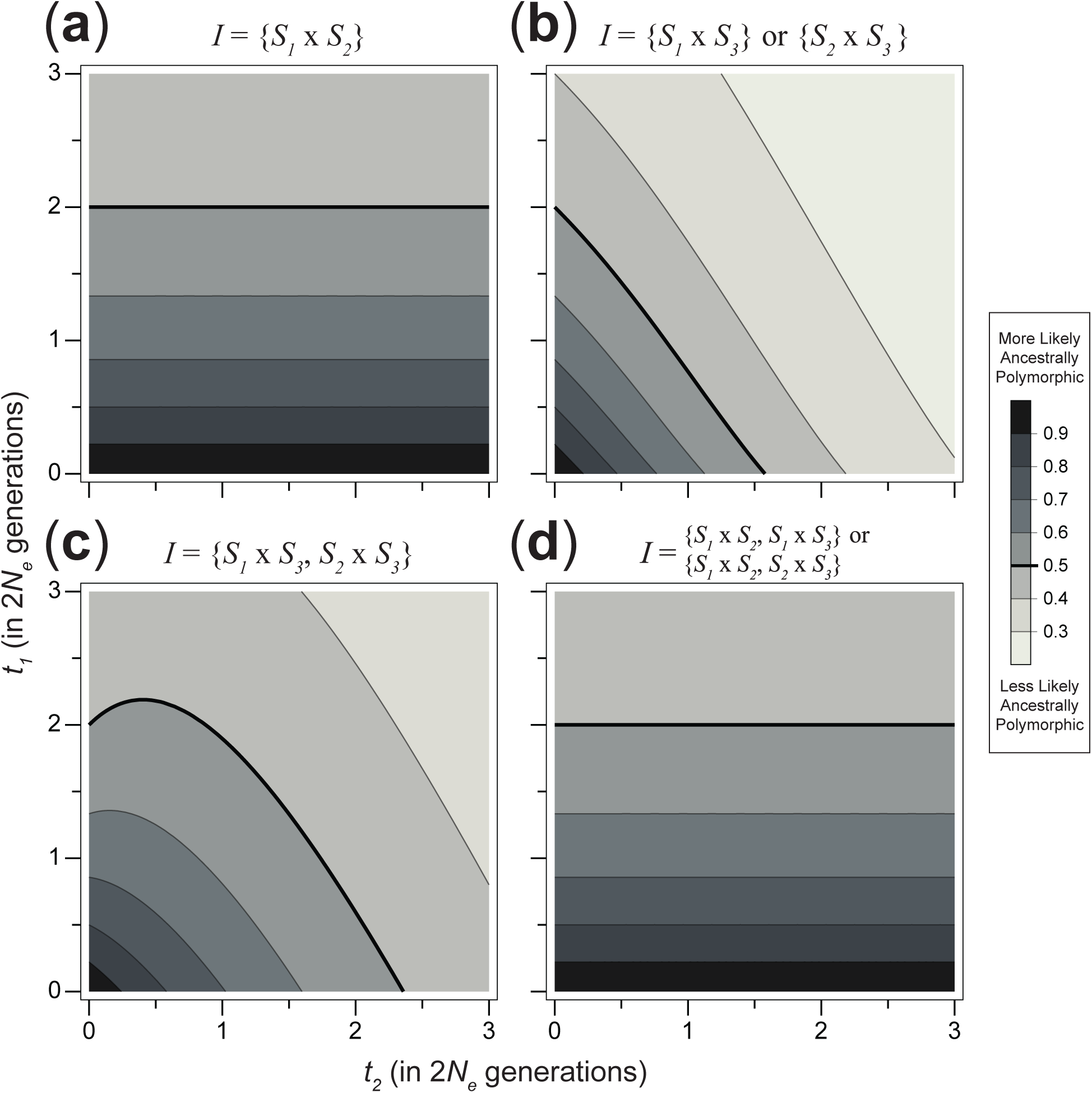
Probability that an incompatibility involves an ancestrally arising locus conditioned on different patterns of reproductive isolation. DMIs are modeled with loci subject to ILS.

**Figure S7.**
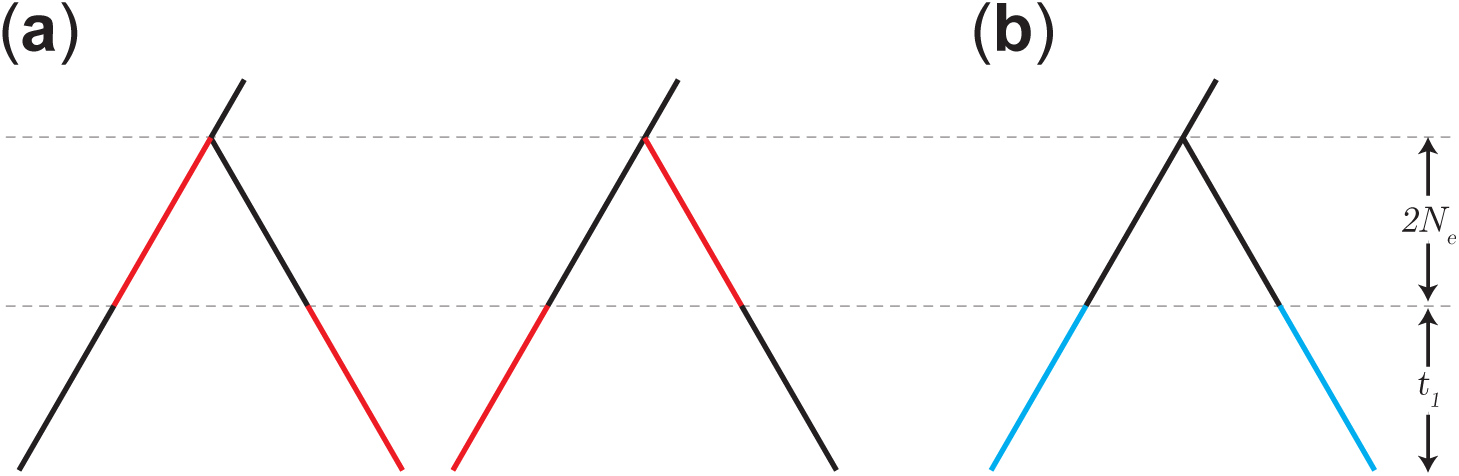
The probability of an ancestrally arising allele in a DMI isolating sister taxa Highlighted segments show the branch segments on which incompatible pairs of alleles can arise. The branch segments on which incompatibilities involving ancestrally arising alleles could arise are shown in (a). The two segments for each of the two panels have lengths, 2*N*_e_, the expected time to coalescence of two lineages, and *t*_1_, the time since divergence to the present, respectively. Branch segments on which incompatibilities from which mutations after divergence produce incompatible alleles are shown in (b), where each of the two segments have length *t_1_*.

**Figure S8.**
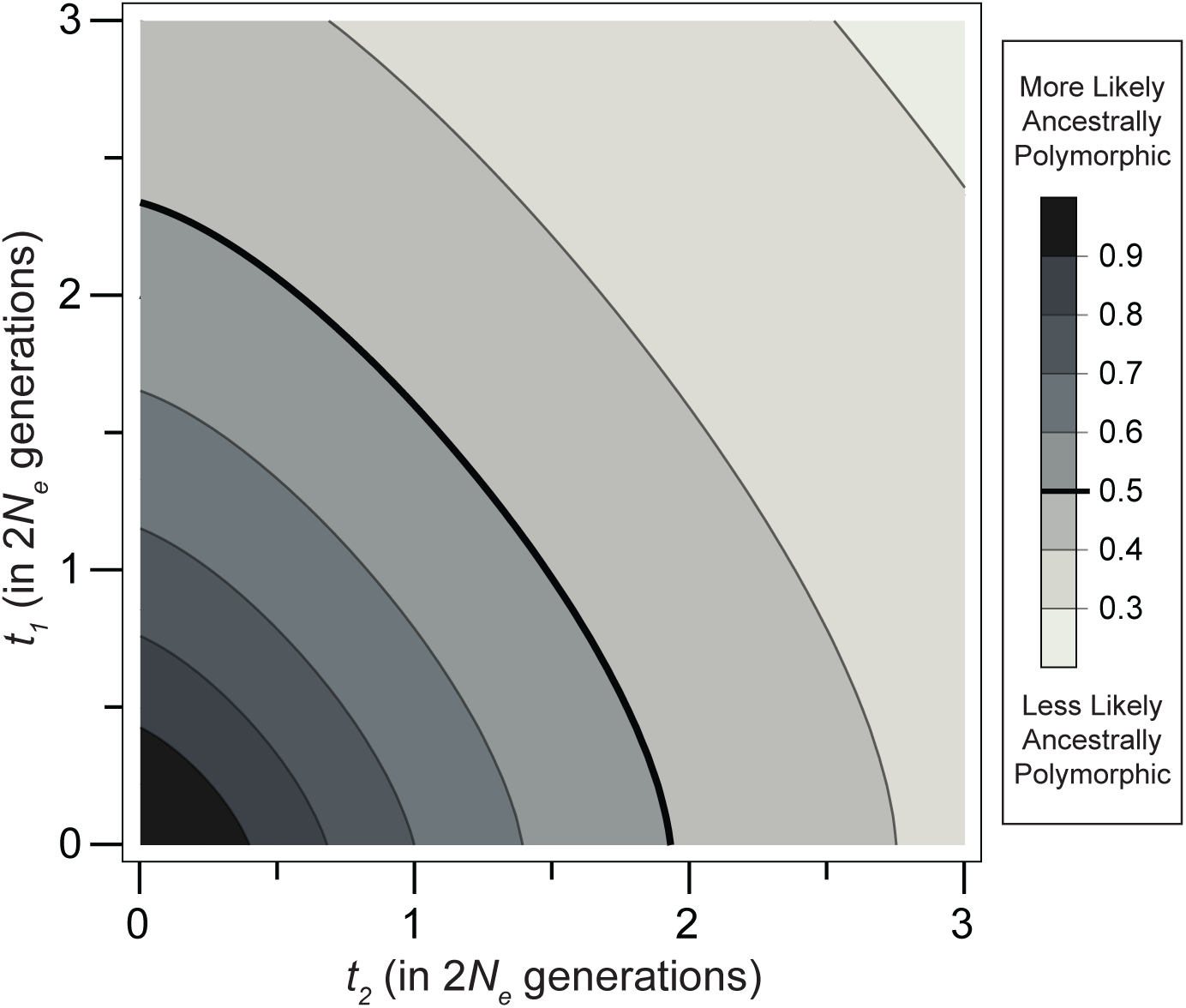
Probability that an incompatibility involves an ancestrally arising locus; model of DMI with potentially polymorphic loci in addition to being subject to ILS. Bold line shows the contour where a pairwise DMI is equally likely to form from at least one ancestrally arising locus as from alleles both derived after divergence.

### Supplemental Methods

#### Probability of a mutation on a branch segment

Assuming mutations are rare, the probability of a mutation on a particular branch segment, i.e. *x_α_* and *y_β_*, is proportional to the length of that segment (Hudson 1992). The probability of a mutation can be estimated as the probability for at least one mutation on the given segment, with no mutations on the rest of the tree,

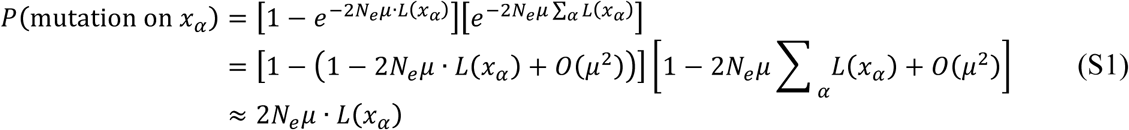
where *2N_eμ_* is the population mutation parameter, *L*(*x_α_*) is the length of branch segment *x_α_*, and a Taylor series approximation has been made to remove terms of order *μ*^2^ and higher.

#### *Mutation order for* (S_1_, S_2_) S_3_ int. *gene trees*

Let *τ_1_* be a random variable representing the time from the *S_1_, S_2_* divergence to the first mutation on segment *g-h* on the first tree. Let *v_1_* and *v_2_* be random variables representing the time from the *S_1_*, *S_2_* divergence to the most recent coalescence on the first and second tree, respectively. The first mutation on segment *g-h* is unconstrained and its probability remains *t_2_* − *q*(*t_2_*). The second mutation on segment *d-g* is constrained by both the timing of the first mutation as well as the timing of the coalescence event at one end of the segment (see Fig. S9). If the mutation on the first tree occurs before the coalescence on the second tree, the second mutation can arise on the entire length of the *d-g* segment, up to time *v_2_*. If the coalescence on the second tree occurs before the first mutation, the second mutation may only arise on the truncated *d-g* segment, up to length *τ_1_*. The probability of an incompatible mutation arising on *d-g* is the expected length,

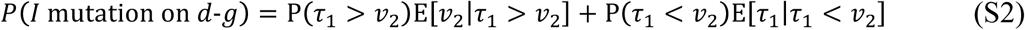

Let *f_v_* be the probability distribution function for *v_1_* and *v_2_* (as described earlier in Equation 11; main text), and *f_τ_* be the probability distribution function for *τ_1_*. Because we assume only a single mutation occurs on segment *g-h*, the probability of *τ_1_* is uniformly distributed along its length, *t_2_* − *v_1_*. The probability distribution function, *f_τ_*, is then

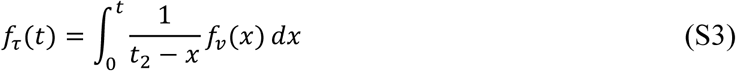

The conditional expectations can thus be expressed as

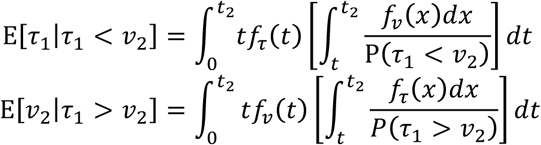

When these expectations are substituted into Equation S2, the probability of an incompatible mutation arising on *d-g* becomes,

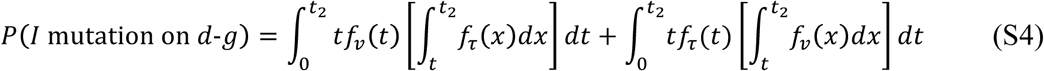

The probability of an incompatibility between these segments can be calculated from the product of the unconstrained mutation probability on segment *g-h, t_2_* − *q*(*t_2_*), and Equation S4. Since segments *d-g* and *g-h* exist on both trees, this product contributes twice to the probability of a derived-ancestral incompatibility when both loci have (*S_1_, S_2_*) *S_3_* int. trees.

**Figure S9.**
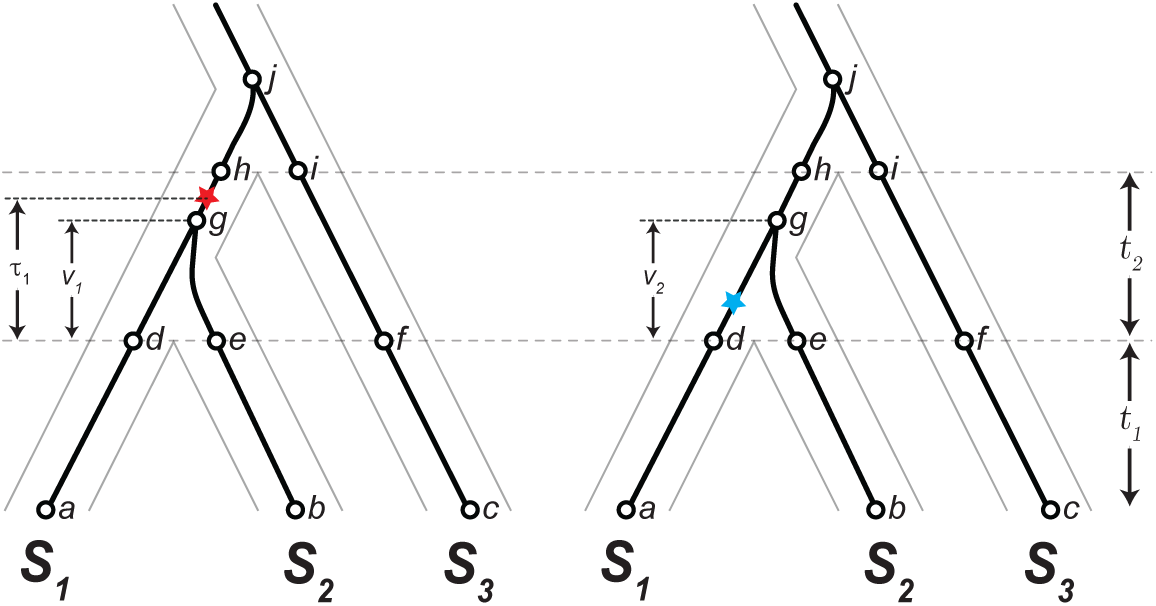
Calculating constraints on derived-ancestral incompatibilities due to mutation order on a pair of (*S_1_*, *S_2_*) *S_3_* int. trees. While this figure shows the mutation on the left tree, marked by the red star, as occurring first, this is not always true. Because *v_1_*, *v_2_*, and *τ_1_* are all random variables, it is possible for the mutation on the second tree, marked by the blue star, to be the first mutation.

#### Mutation order for incompatibilities that arise ancestrally

As previously noted, there are a number of branch segments with ambiguous endpoints in the ancestral population, forcing us to consider mutation order for derived-ancestral incompatibilities when incompatible alleles are allowed to arise before divergence. These calculations follow Eqs. S2 and S4; the timing of the first mutation restricts the length of the segment on which the second mutation may occur. Let *λ_Ai_* be a random variable representing the length of the segment on which an unrestricted second mutation might occur for a pattern of branch segment asymmetry, *A_i_*. Let *τ_Ai_* be a random variable representing the time from the oldest divergence to the first mutation for this asymmetry pattern. The probability of the first mutation is unrestricted and equal to the expected length of the first branch segment. The probability of the second mutation depends on whether the first mutation precedes the coalescence of the second branch segment and can be expressed as

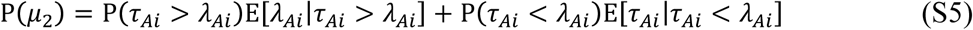

Branch segments with mutation order asymmetry affecting the probability of derived-ancestral incompatibilities are labeled with their respective asymmetry patterns in Appendix 2. The probability distribution functions for *λ_Ai_* can be calculated from the time to coalescence for three-to-two and two-to-one lineages. Similarly, *τ_Ai_* can be calculated from the convolution of these functions and the probability distribution function for a mutation uniformly distributed along the appropriate branch. The probability distribution functions for *λ_Ai_* and *τ_Αi_*, as well as Ρ(*μ_1_*) and Ρ(*μ_2_*), are listed for each asymmetry pattern in Table S1.

We modify Equation 13 (main text) to calculate the probability of concordance with polymorphic incompatibilities. Let **1R***_Tx,Ty_* be the indicator matrix for segments on trees *T_x_* and *Ty*, whose elements are 1 when the segment combination can lead to an incompatibility and 0 otherwise. Let **Α***_Tx,Ty_* be a matrix whose elements are the probabilities for a derived-ancestral incompatibility that depends on mutation order (i.e. Ρ(*μ_1_*) · Ρ(*μ_2_*) for that segment combination). The probability of an incompatibility with isolation pattern *I*, conditioned on gene trees *T_x_* and *T_y_*, is then

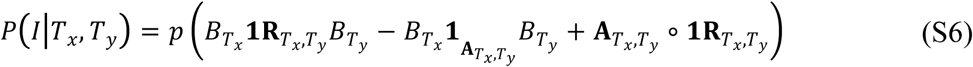
where 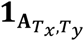 is a matrix whose elements are 1 for non-zero entries of **Α***_Tx,Ty_* and 0 otherwise.

As before, we can form the 4x4 matrix, ***D***(*I*), for each isolation pattern *I* from Equation S6, and calculate concordance probabilities from Equation 4 (main text).

**Table S1.** Probabilities for incompatibility-participating mutations that depend on mutation order Note that patterns A_2_ and A_5_ correspond to calculations from Eqs. 16 (main text) and S4 respectively.

| Pattern | $f_{\lambda_{Ai}}(t)$ | $f_{\tau_{Ai}}(t)$ | $P(\mu_1)$ | $P(\mu_2)$ |
| --- | --- | --- | --- | --- |
| A<sub>1</sub> | $e^{-t}$ | $\int_0^t 3e^{-3(t-s)} \left[ \int_s^\infty \frac{e^{-x}}{x} dx \right] ds$ | 1 | $1 - \frac{3}{4} \log(2)$ |
| A<sub>2</sub> | 1 for $t = t_2$ | $\int_0^t \frac{1}{t_2 - x} f_v(x) dx$ | $t_2 - q(t_2)$ | $\frac{1}{2}(t_2 + q(t_2))$ |
| A<sub>3</sub> | $\frac{3}{2}(e^{-t} - e^{-3t})$ | $\int_t^\infty \frac{e^{-x}}{x} dx$ | 1 | $\frac{4}{3} - \frac{25}{18} \log(2)$ |
| A<sub>4</sub> | $3e^{-3t}$ | $\int_t^\infty \frac{e^{-x}}{x} dx$ | 1 | $\frac{1}{3} - \frac{1}{9} \log(4)$ |
| A<sub>5</sub> | $f_v(t)$ | $\int_0^t \frac{1}{t_2 - x} f_v(x) dx$ | $t_2 - q(t_2)$ | (see Eq. S4) |
| A<sub>6</sub> | $3e^{-3t}$ | $\int_0^t 3e^{-3(t-s)} \left[ \int_s^\infty \frac{e^{-x}}{x} dx \right] ds$ | 1 | $\frac{1}{3} - \frac{1}{9} \log(2)$ |
| A<sub>7</sub> | $\frac{3}{2}(e^{-t} - e^{-3t})$ | $\int_0^t 3e^{-3(t-s)} \left[ \int_s^\infty \frac{e^{-x}}{x} dx \right] ds$ | 1 | $\frac{4}{3} - \frac{77}{72} \log(2)$ |

## Appendix 1. Incompatibility Matrices

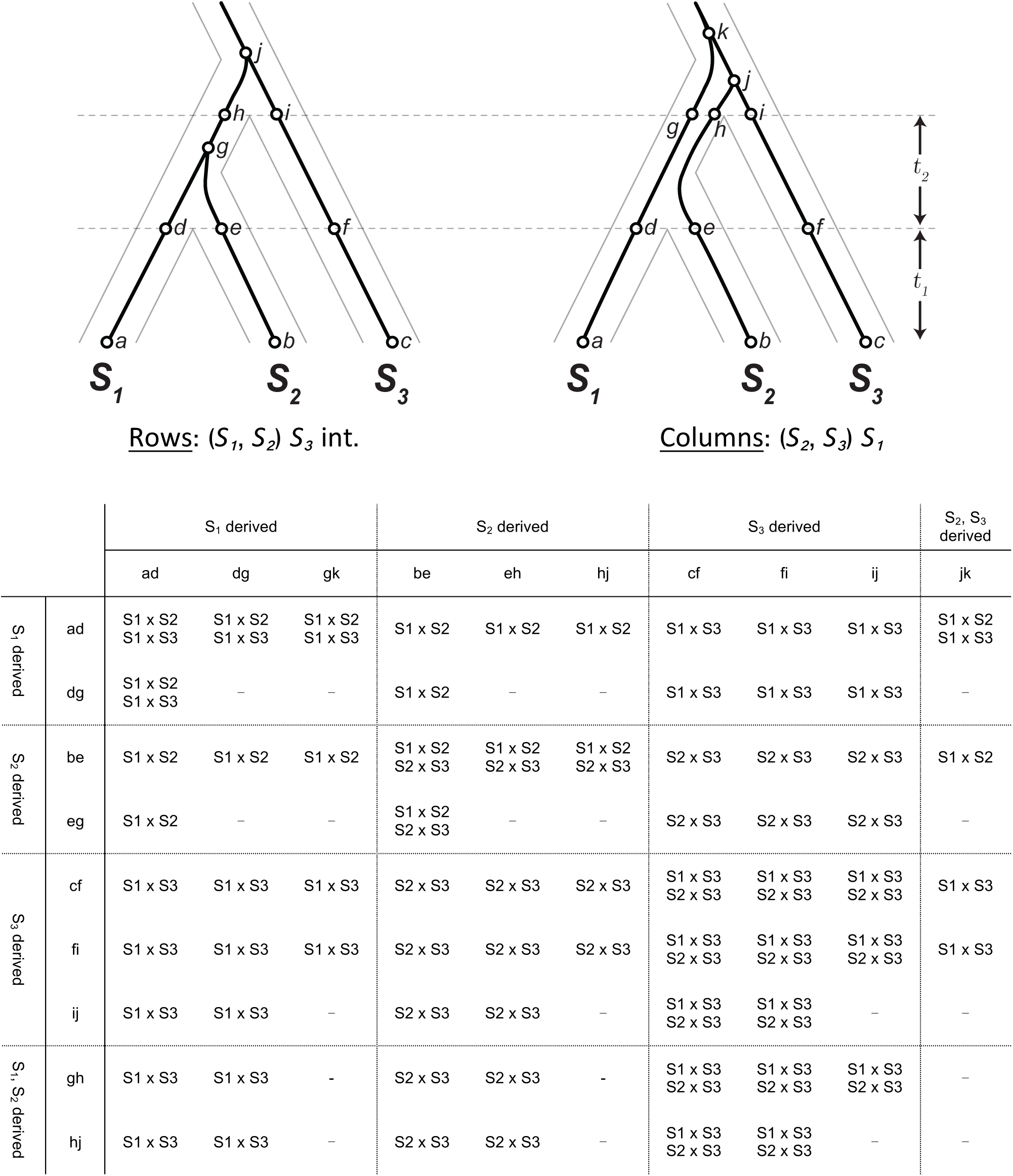
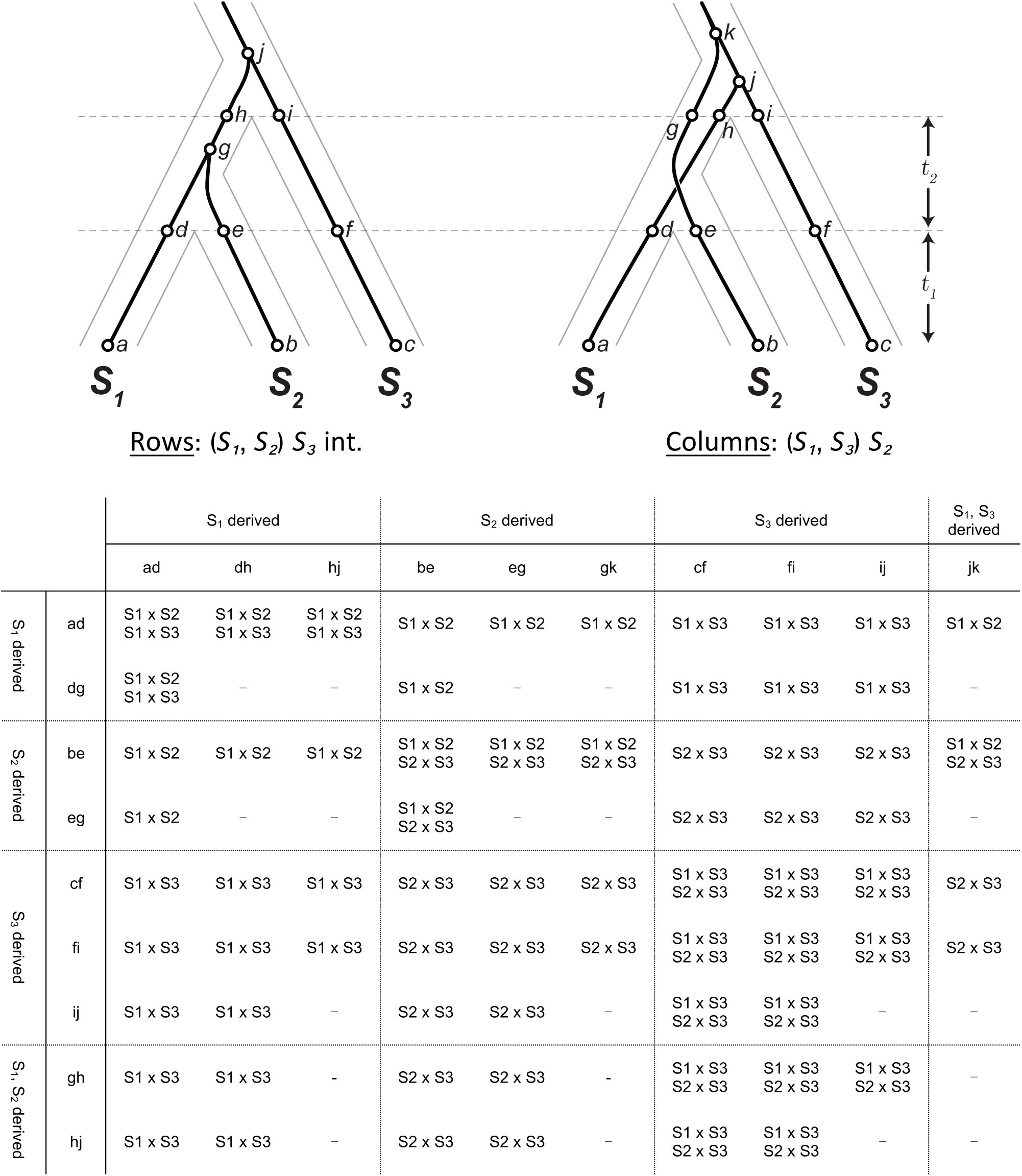
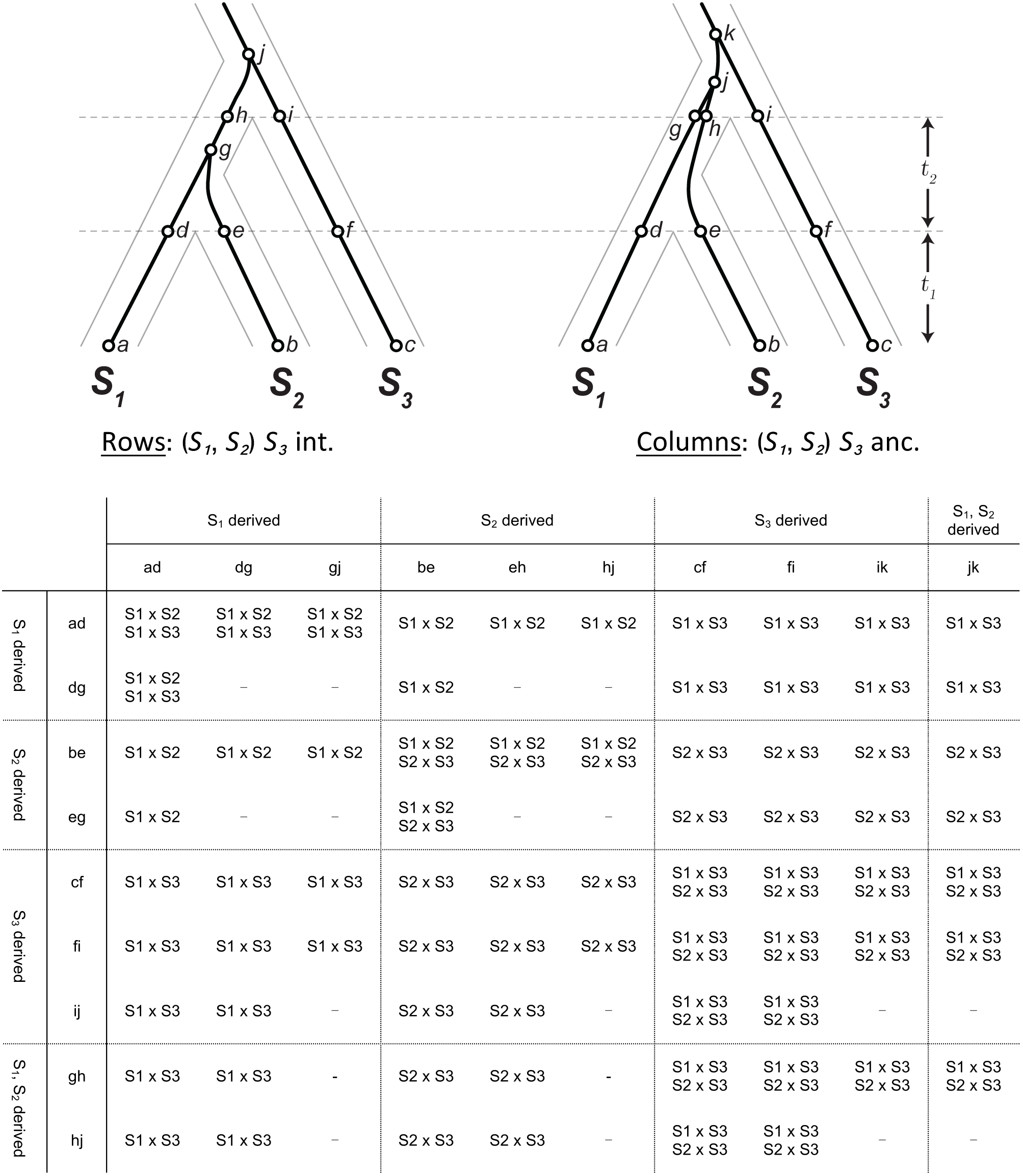
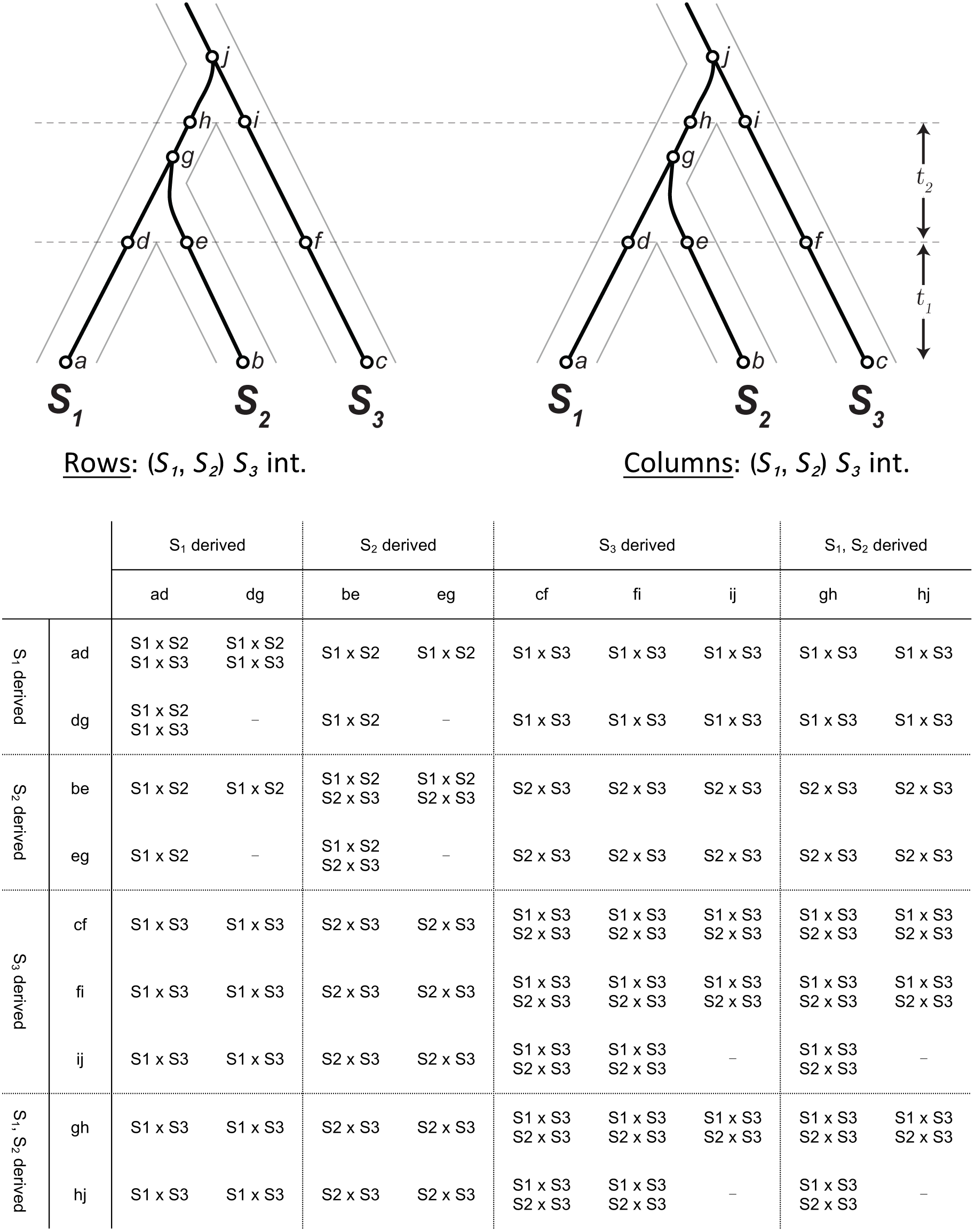
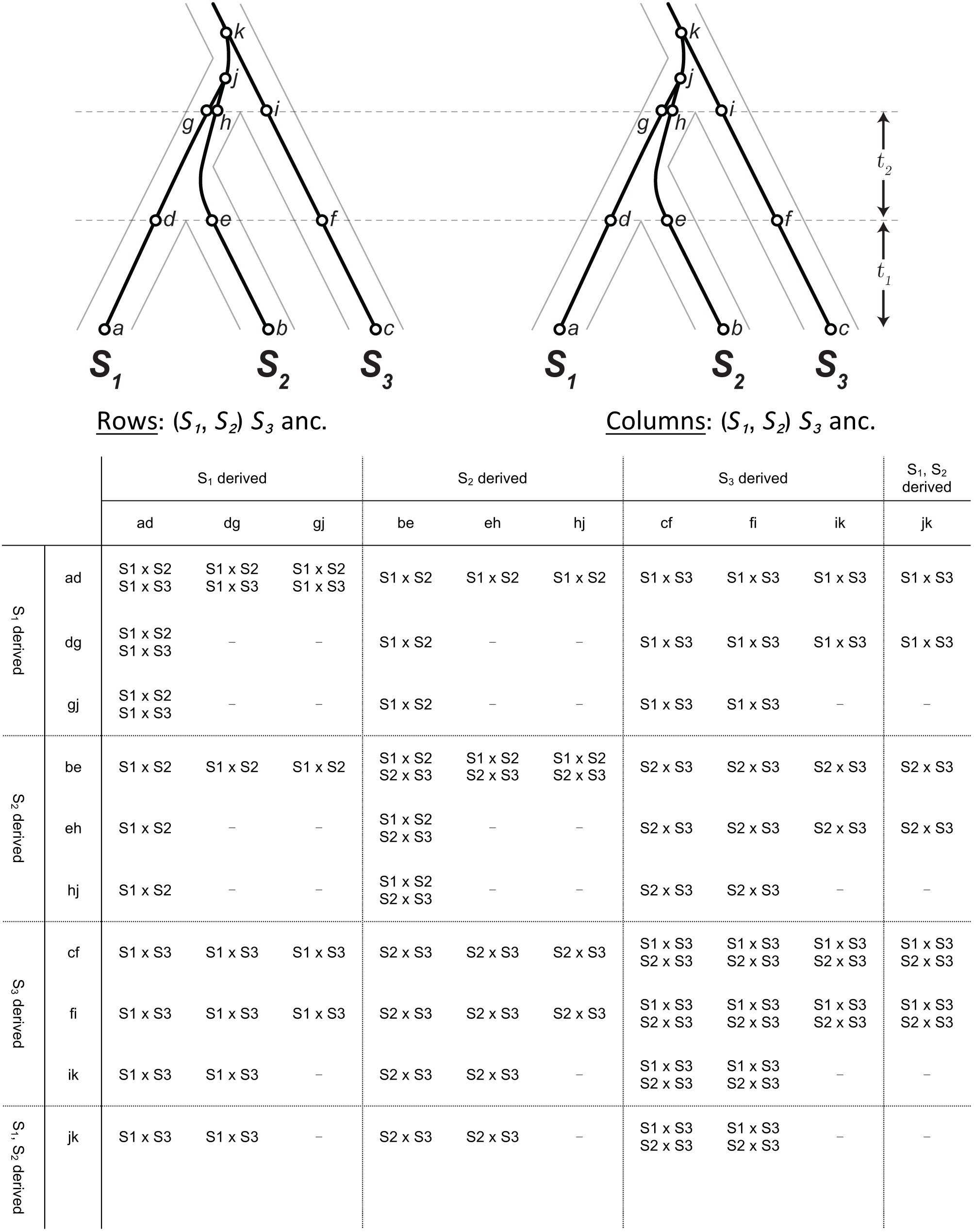
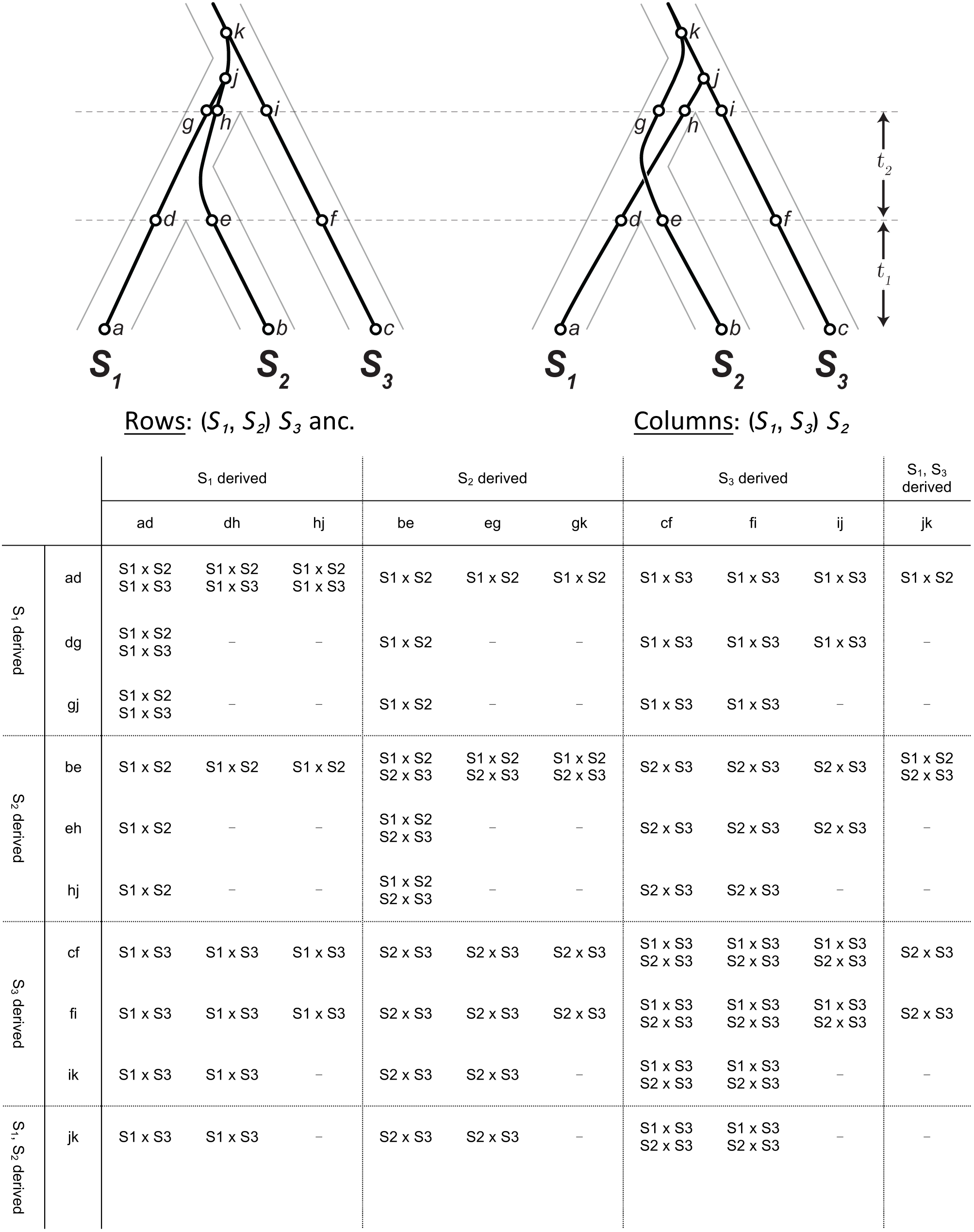
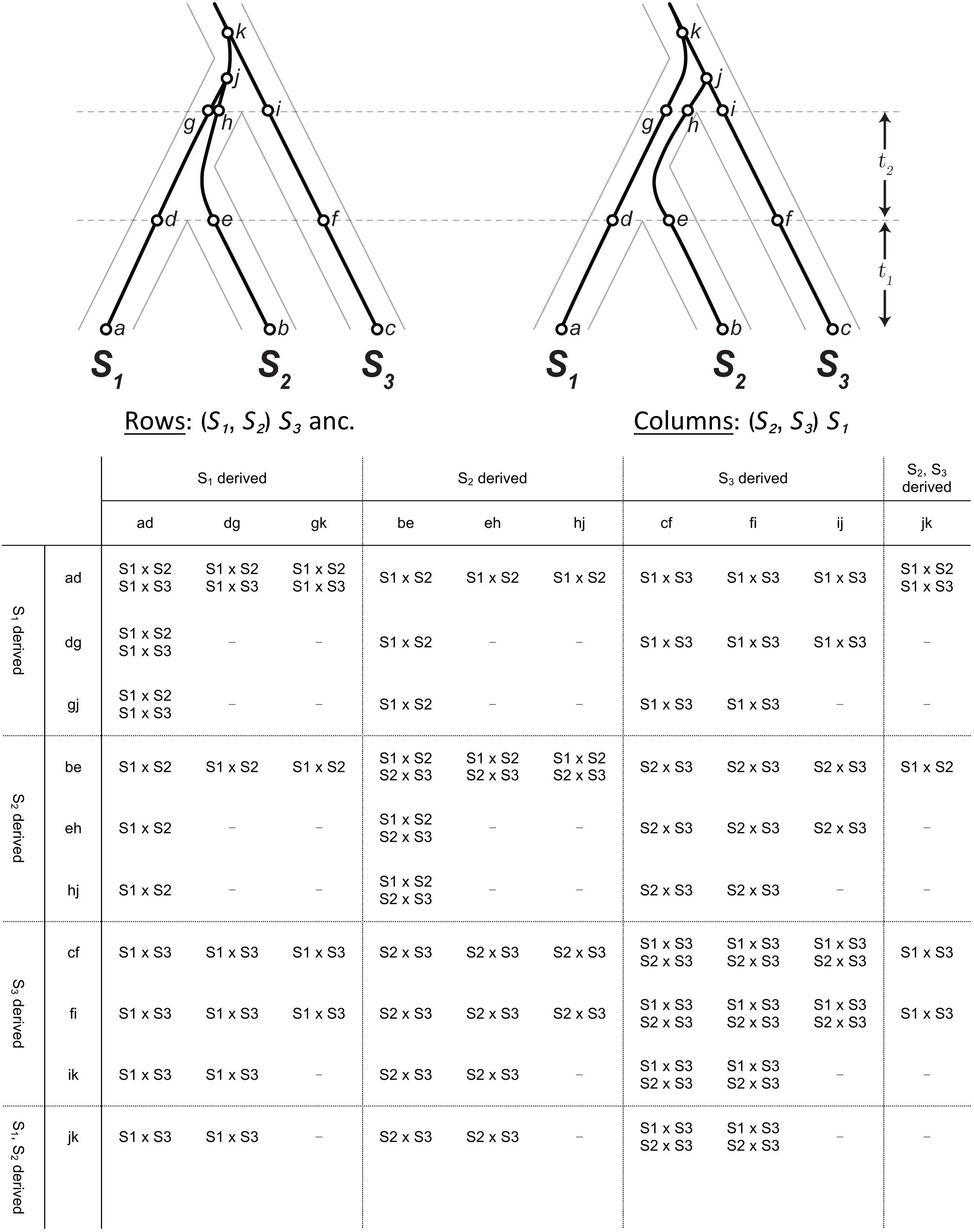
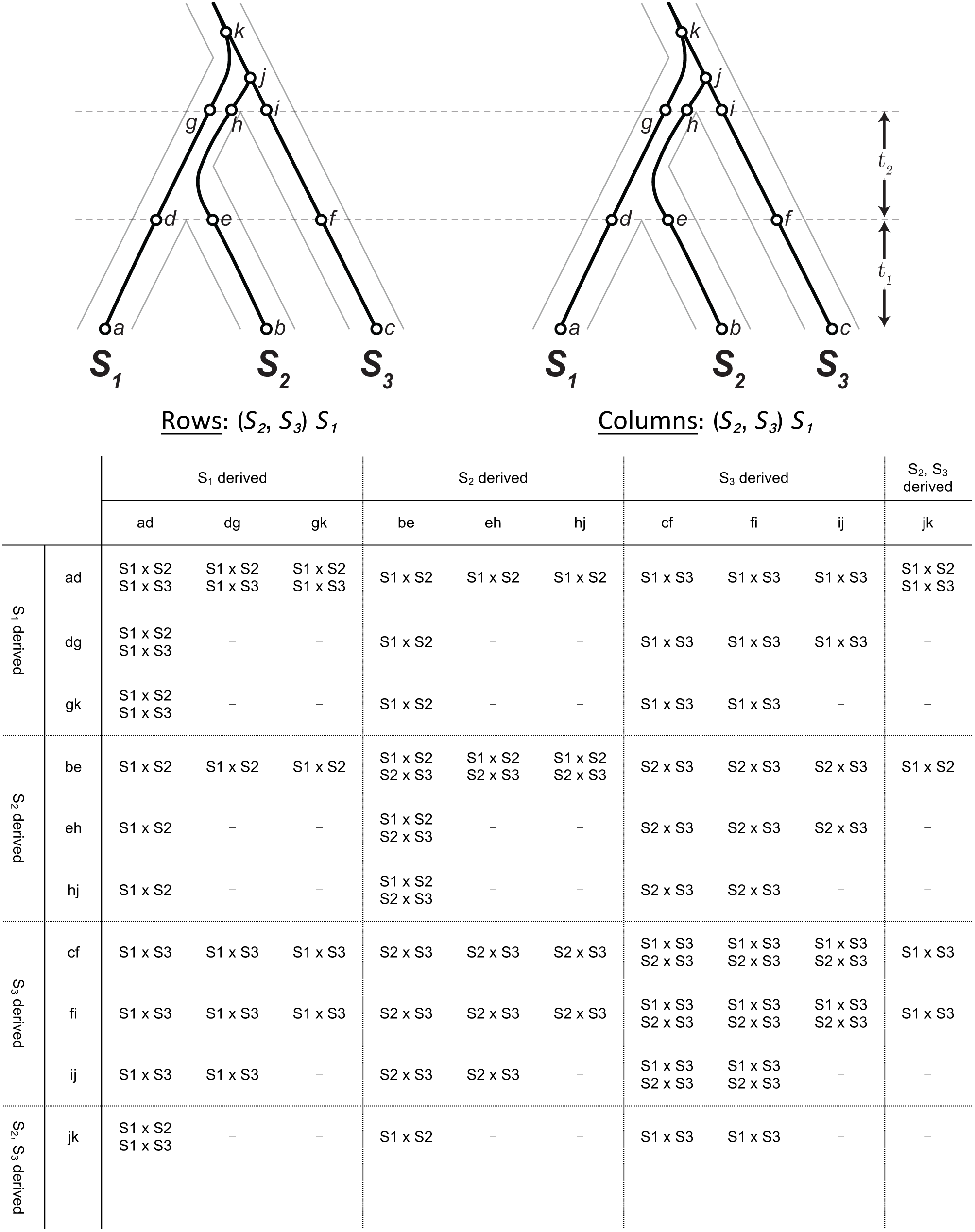
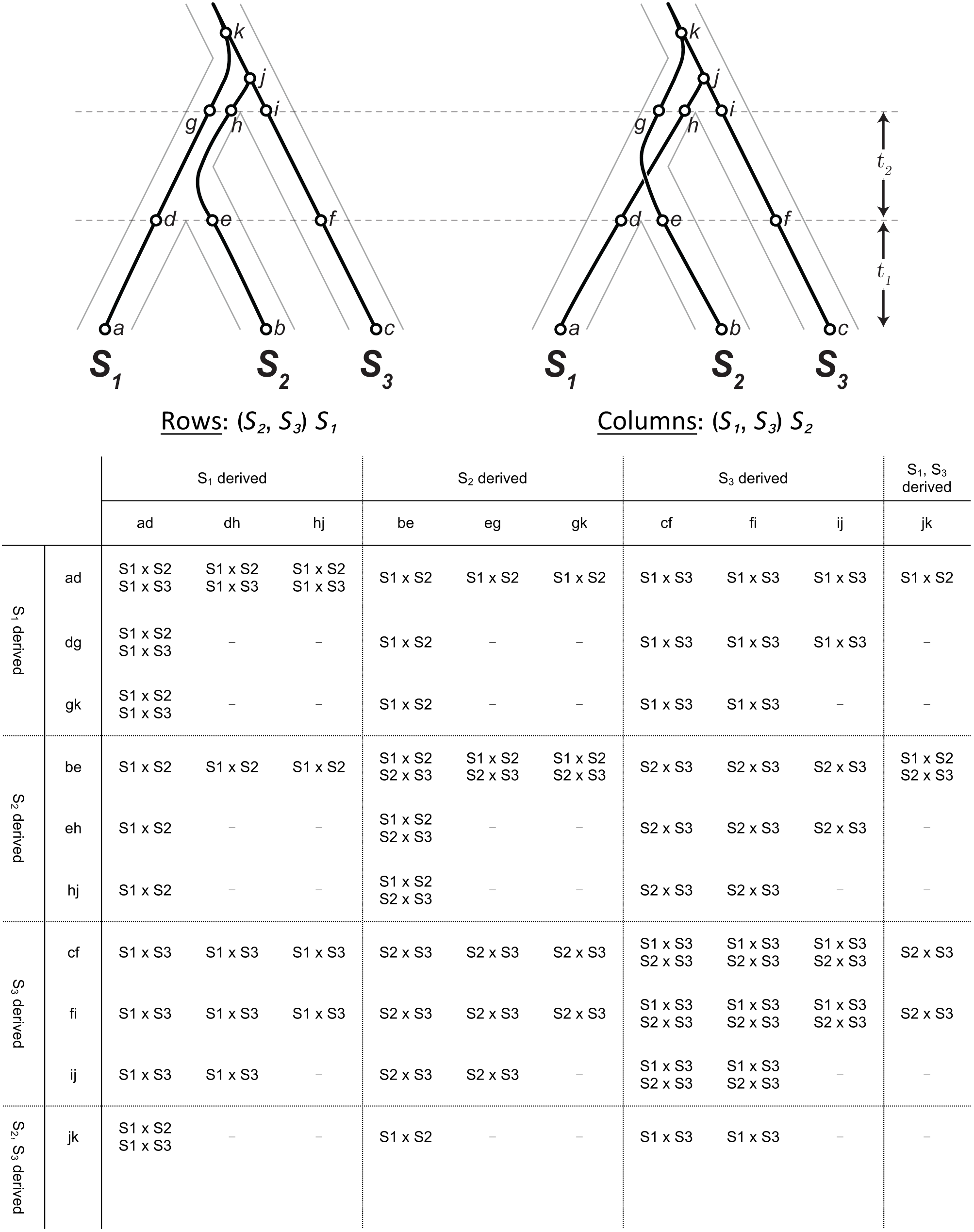
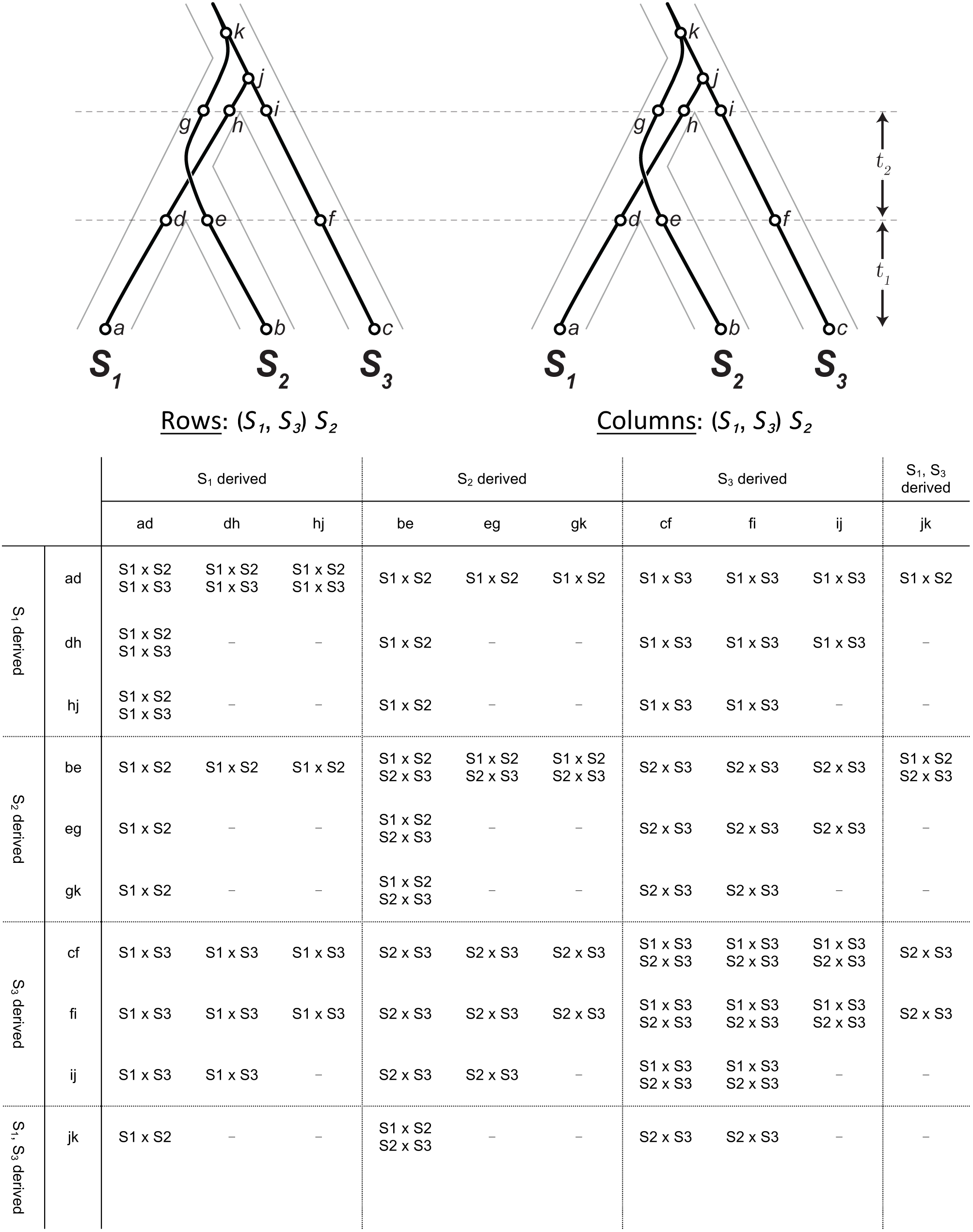

## Appendix 2. Incompatibility Matrices (polymorphic incompatibilities allowed)

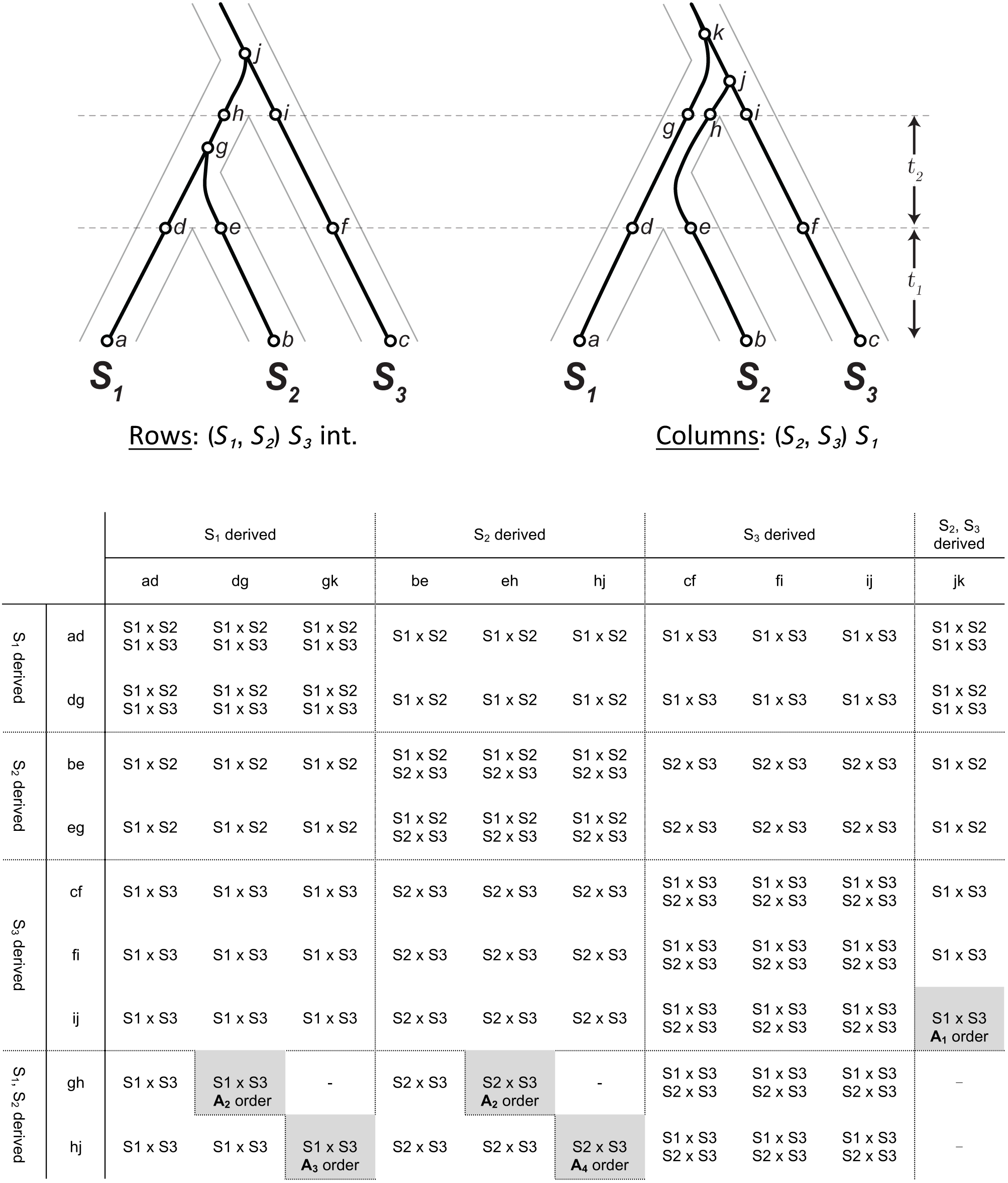
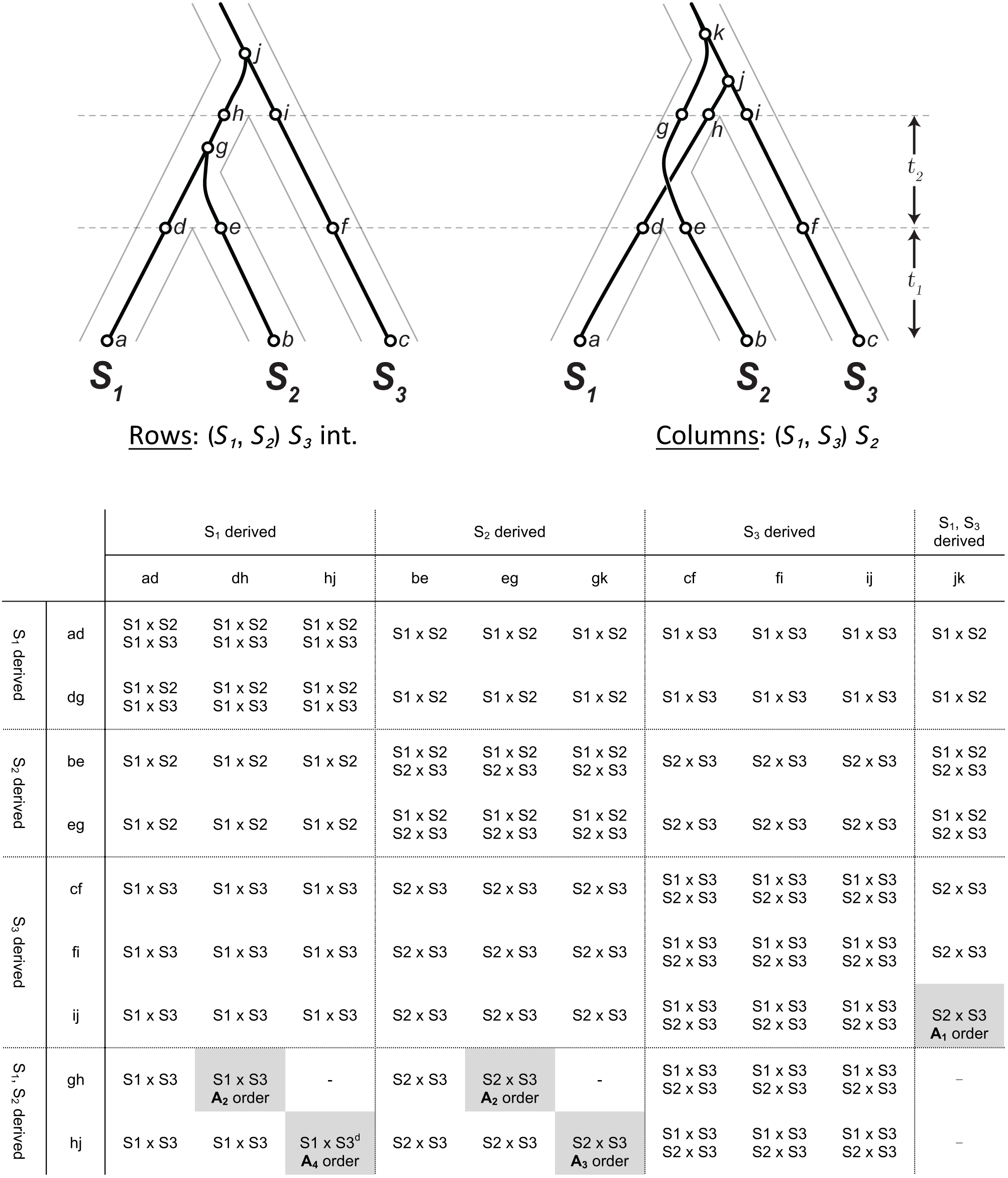
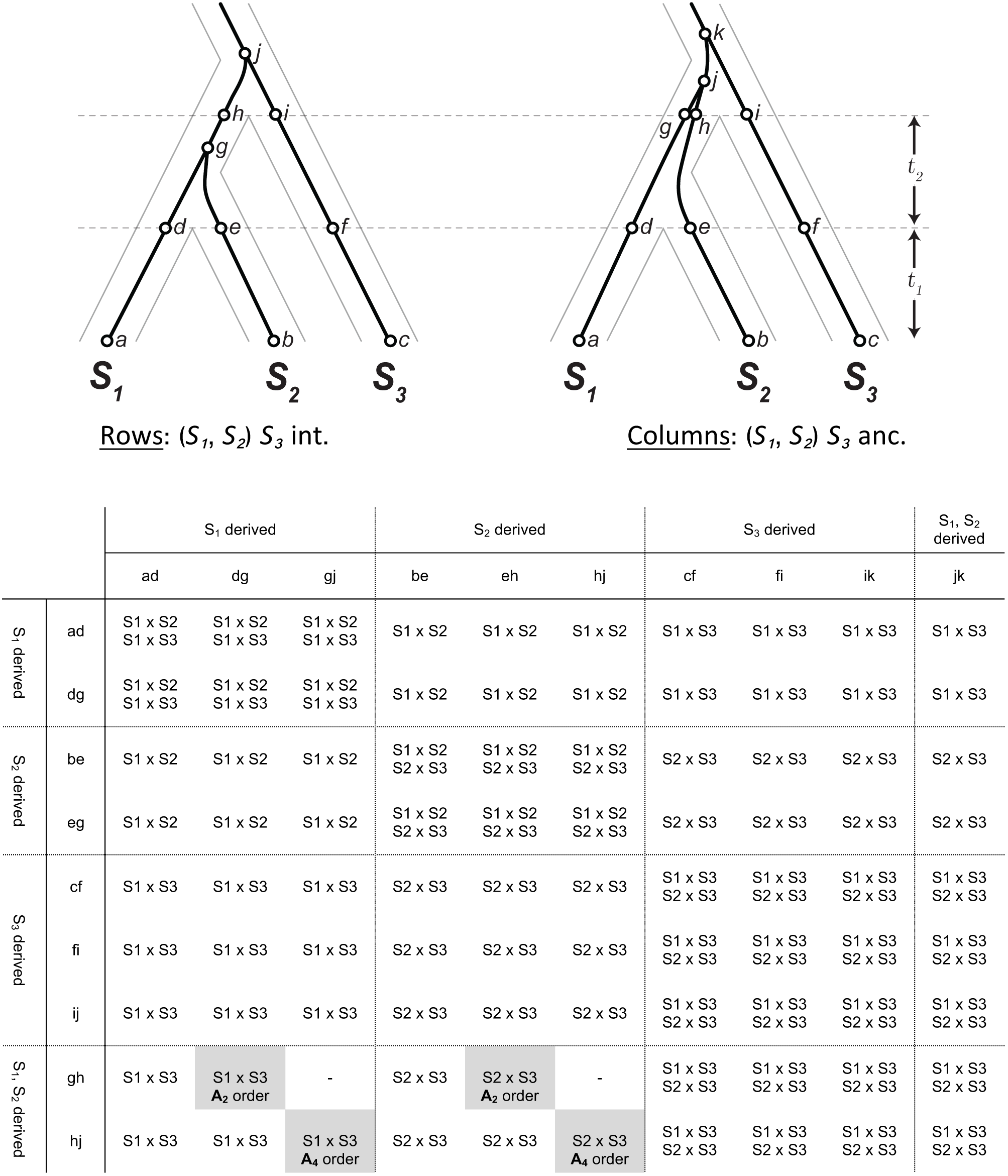
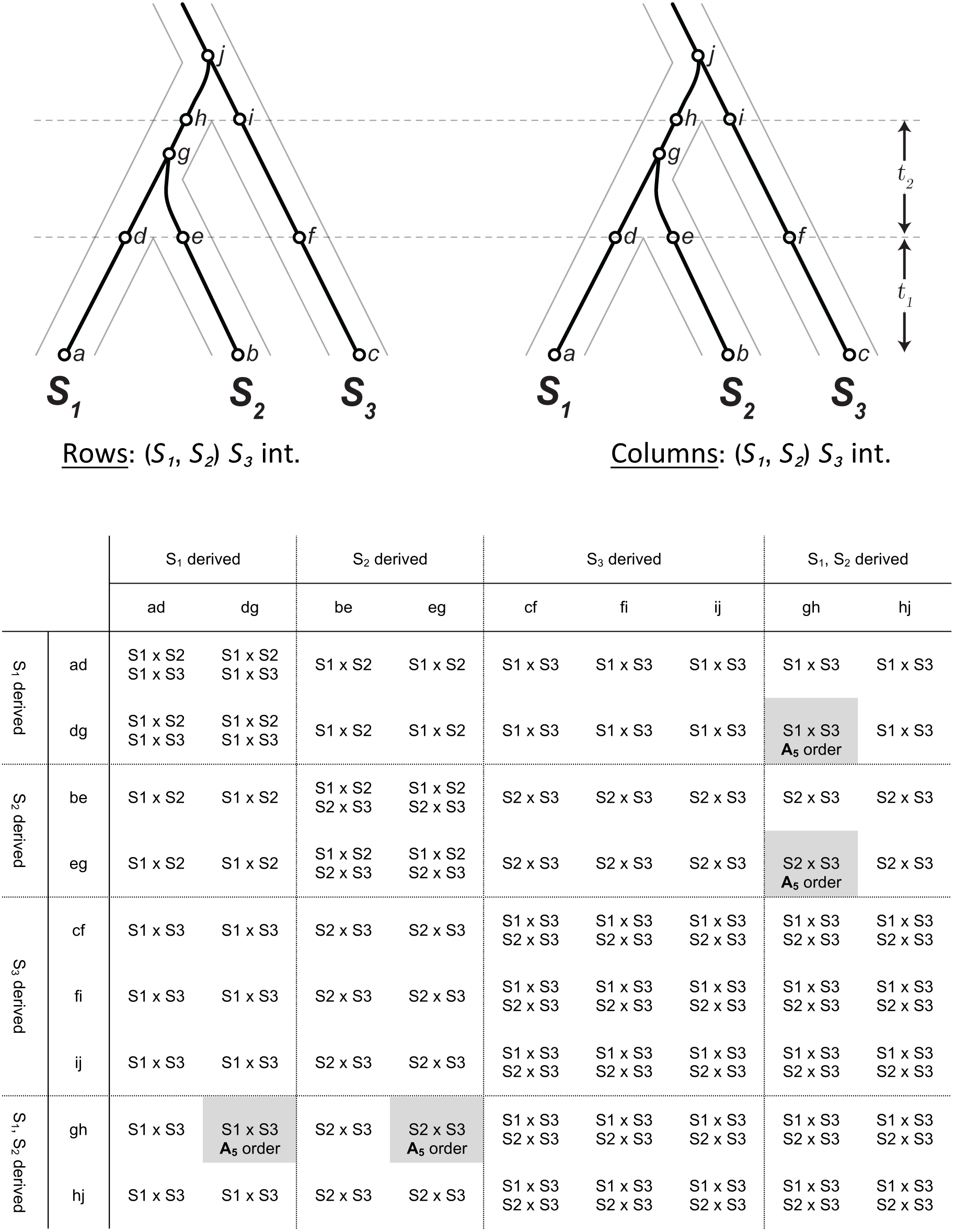
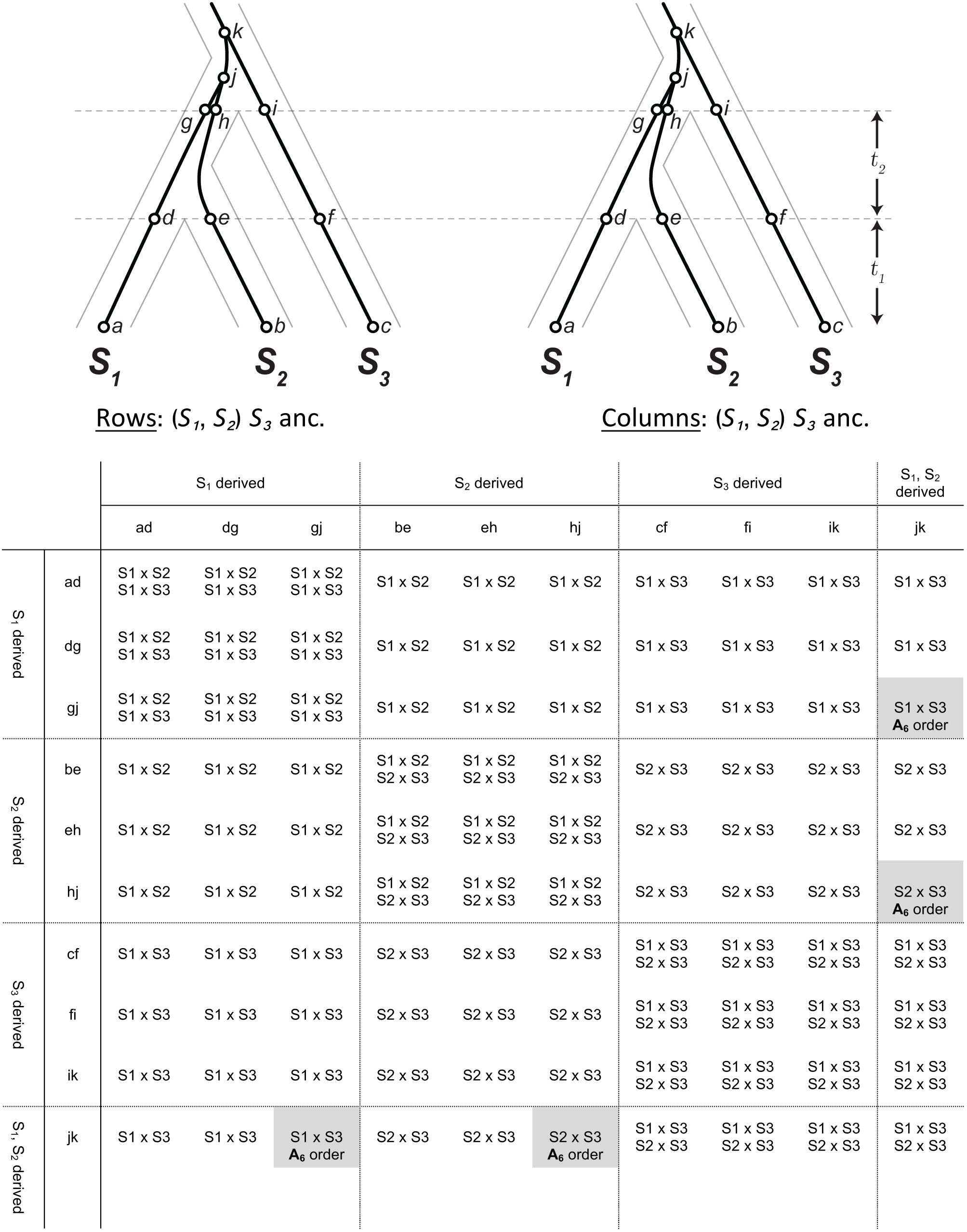
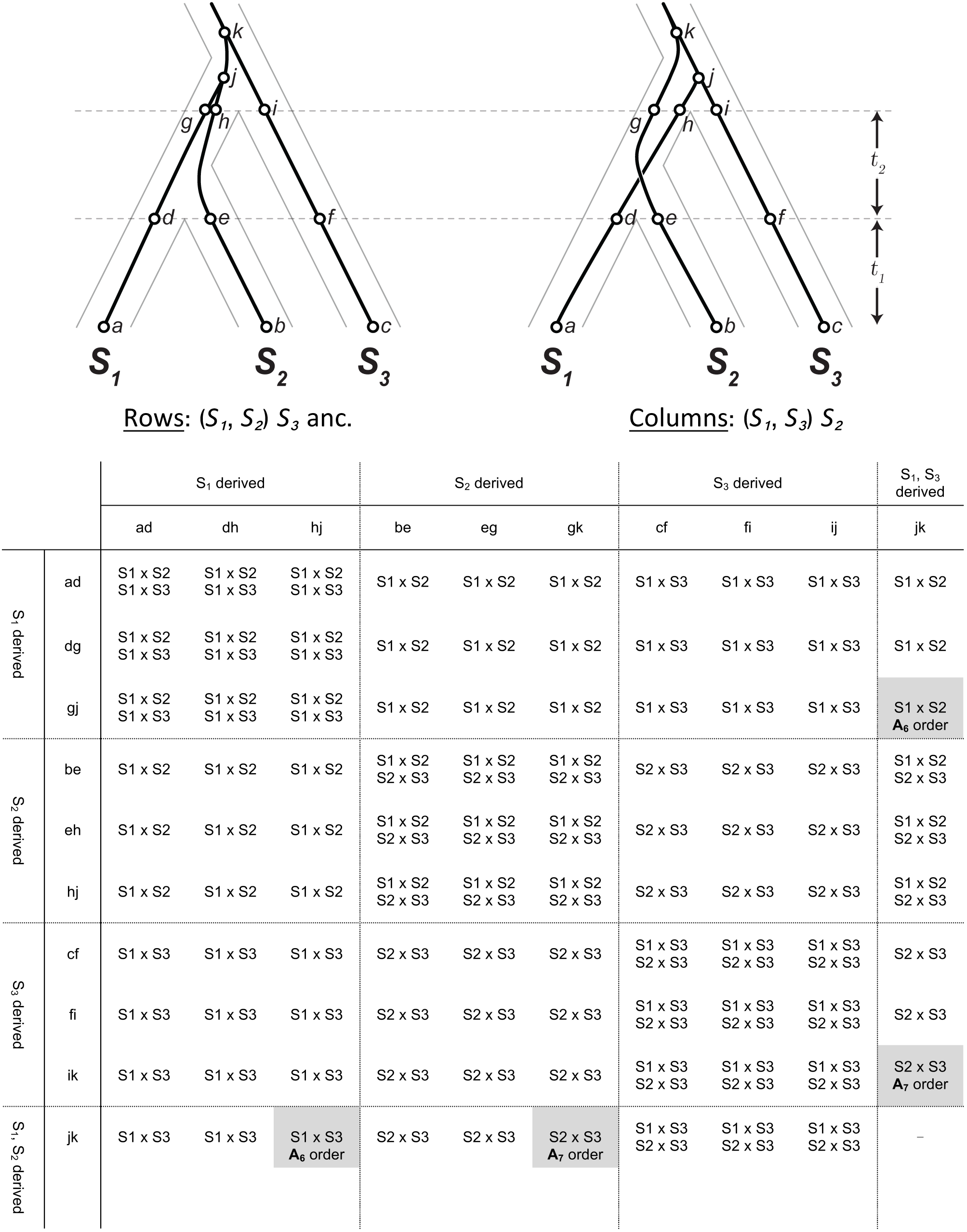
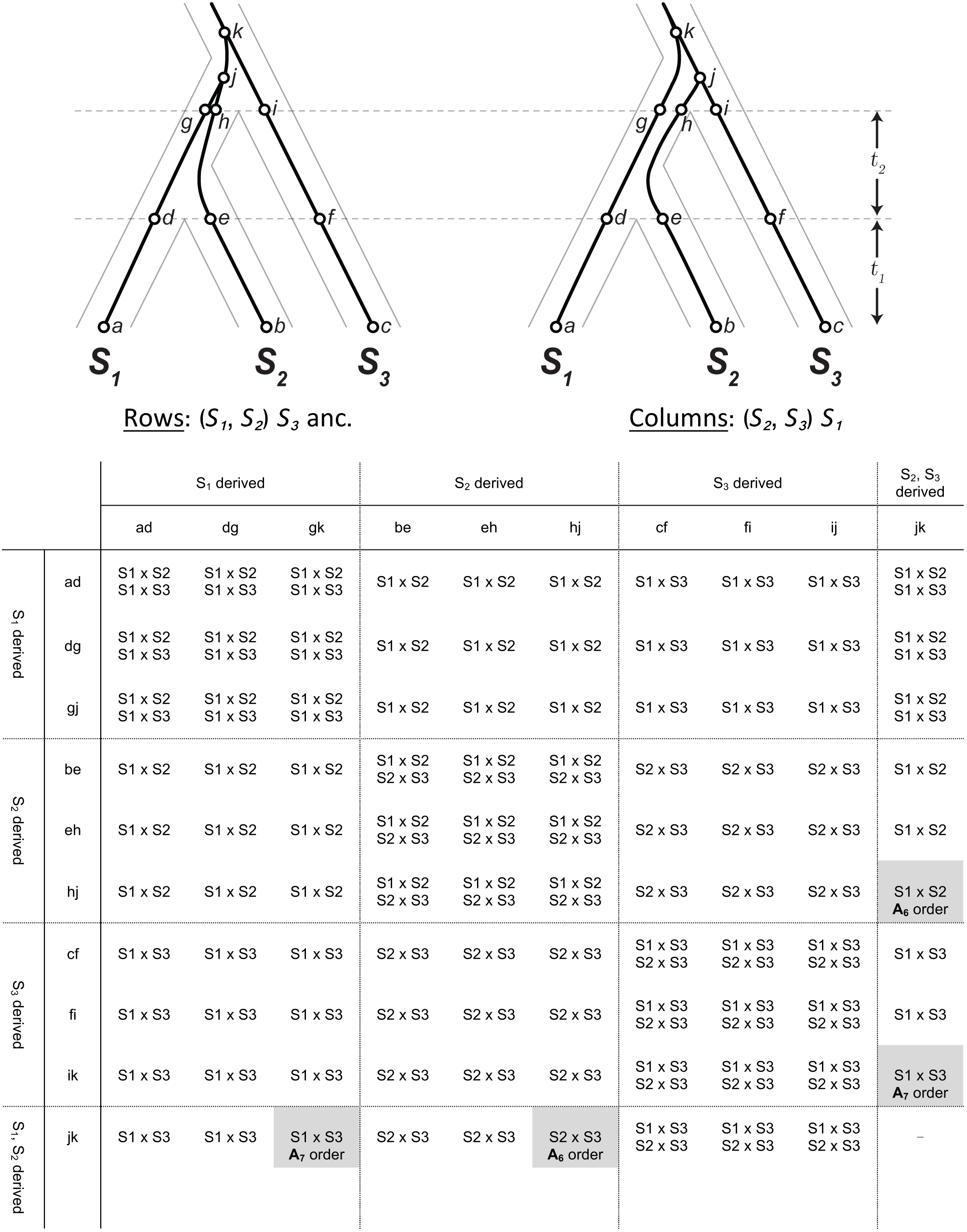
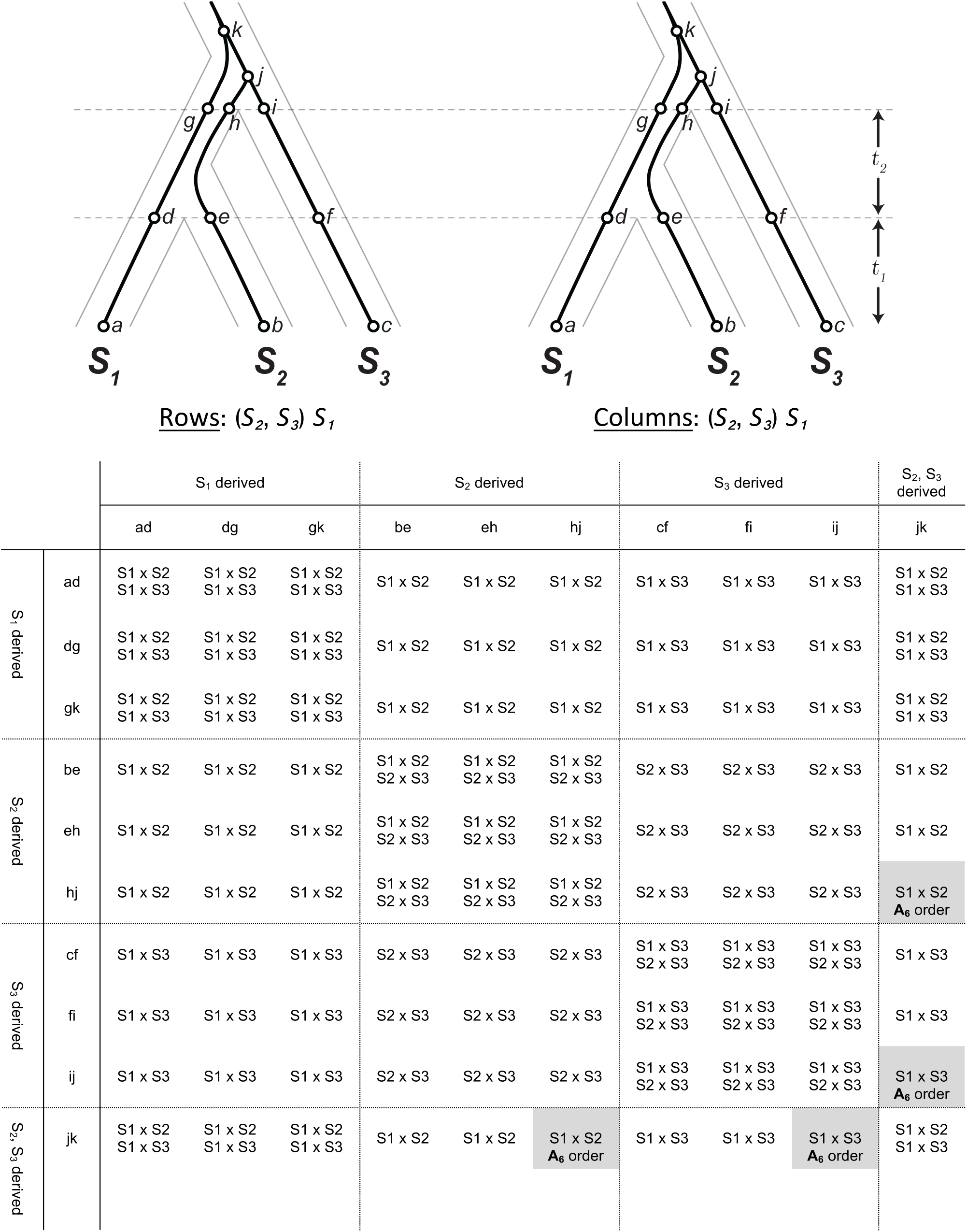
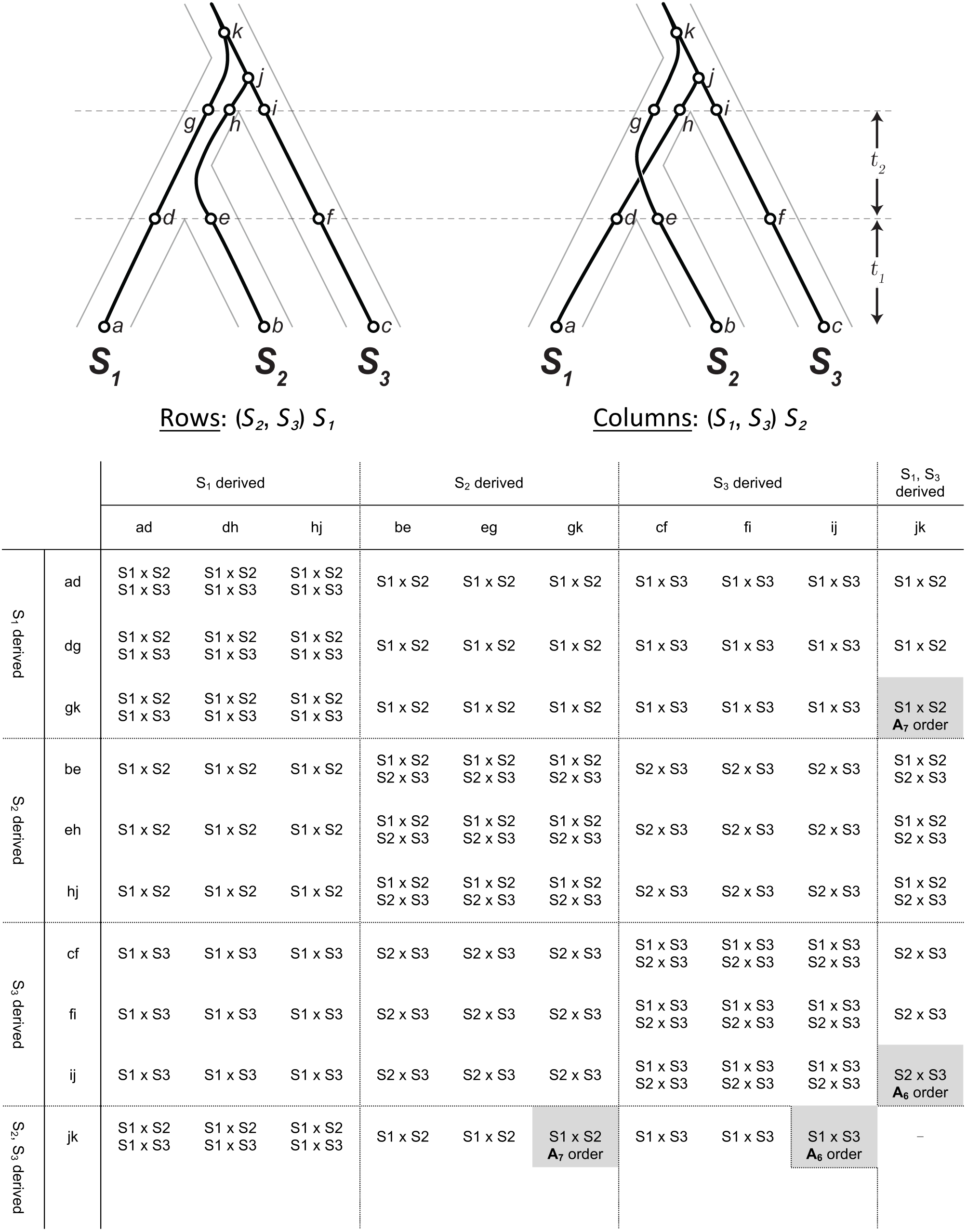
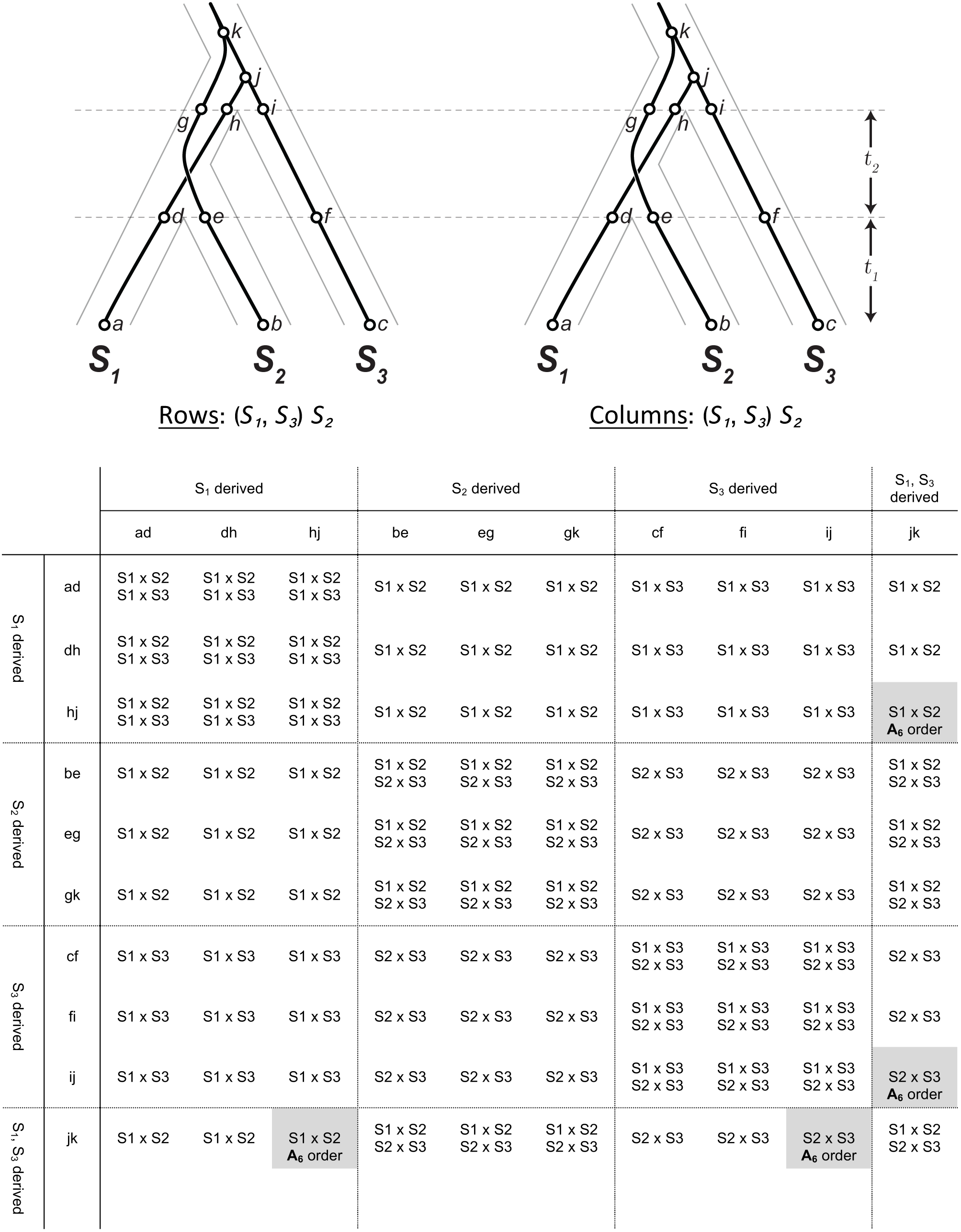

